# Jointly Defining Cell Types from Multiple Single-Cell Datasets Using LIGER

**DOI:** 10.1101/2020.04.07.029546

**Authors:** Jialin Liu, Chao Gao, Joshua Sodicoff, Velina Kozareva, Evan Z. Macosko, Joshua D. Welch

## Abstract

High-throughput single-cell sequencing technologies hold tremendous potential for defining cell types in an unbiased fashion using gene expression and epigenomic state. A key challenge in realizing this potential is integrating single-cell datasets from multiple protocols, biological contexts, and data modalities into a joint definition of cellular identity. We previously developed an approach called Linked Inference of Genomic Experimental Relationships (LIGER) that uses integrative nonnegative matrix factorization to address this challenge. Here, we provide a step-by-step protocol for using LIGER to jointly define cell types from multiple single-cell datasets. The main steps of the protocol include data preprocessing and normalization, joint factorization, quantile normalization and joint clustering, and visualization. We describe how to jointly define cell types from single-cell RNA-seq and single-nucleus ATAC-seq data, but similar steps apply across a wide range of other settings and data types, including cross-species analysis, single-nucleus DNA methylation, and spatial transcriptomics. Our protocol contains examples of expected results, describes common pitfalls, and relies only on our freely available, open-source R implementation of LIGER. We also provide Rmarkdown tutorials showing the outputs from each individual code segment. The analysis process can be performed in 1 - 4 h depending on dataset size and assumes no specialized bioinformatics training.

## Introduction

Identifying the molecular features that define the types and functions of individual cells provides a tremendous opportunity for understanding the genomic blueprint of the human body. The classic approach to categorize cells relies on qualitative characterization, including gross morphology, the presence or absence of a few surface proteins, and broad cellular function. However, a more comprehensive definition of cell identity requires the inclusion of transcriptomic and epigenomic profiles of cells. In recent years, a variety of high-throughput single-cell sequencing technologies have emerged, measuring the gene expression, DNA methylation, and chromatin accessibility of individual cells. These data modalities together enable researchers to revisit the conventional classifications of cell types and states in a quantitative, systematic, unbiased fashion. Such quantitative definition of cell identity promises to revolutionize our understanding of cell biology across a range of contexts, including neuroscience and developmental biology. A reference map of the molecular states of healthy cells will in turn allow for probing the causes of cellular abnormality and may ultimately inspire the development of novel targeted therapeutics.

In order to achieve this goal, an analytical method capable of integrating various single-cell data modalities is needed. Although large datasets of expression, DNA methylation, chromatin accessibility at single-cell resolution are widely available, multiple modalities are not usually measured from the same individual cells due to limitations in existing sequencing technologies. This requires the integration method to identify the features or properties that represent the “essential” aspects of a cell’s identity, rather than the “dispensable” properties that change across biological settings, modalities, protocols, or time.

In addition, when leveraging such datasets to define cell identity, we want to capture both discrete cell types and continuous variation such as cell states. For example, Saunders et al. found that glutamatergic and GABAergic neurons in the mouse cortex specialize in clearly distinguishable, discrete subtypes^1^. In contrast, the spiny projection neurons in the striatum show more continuous variation, with cell identity being the combination of patch/matrix and direct/indirect distinctions^1^.

Another important consideration is the ability to separate technical confounders from biological signals. Such confounding effects can include the presence of artificial cell doublets created during the cell isolation process, differences in mitochondrial RNA and ribosomal protein content due to cell dissociation, and the presence of free-floating RNA from lysed cells. Failure to account for such factors can lead, for example, to erroneously defining a cell type or state predominantly by its mitochondrial RNA profile, in the absence of any significant biological difference from other cells of the same type.

Integrative analysis should also allow for identifying similarities and differences in corresponding cells across tissues, species, and conditions. For example, it will help to answer questions such as how one tissue differs from another in terms of cell type composition as well as cell-type-specific gene expression. We can also gain a deeper understanding of the cell types and cell-type-specific differences underlying diverse forms and functions across species. Moreover, biomedical researchers are often interested in the cell-type-specific gene expression patterns associated with risk, onset and progression of diseases.

### Development of the protocol

In our previous Cell paper^2^, we first used LIGER to jointly define cell types and their sex-specific differences in the mouse bed nucleus of the stria terminalis (BNST). Through the analysis of scRNA-seq data from BNST, we found significant sexual dimorphism in the gene expression patterns of multiple cell types. Secondly, we utilized LIGER to define cell types across species in mouse and human substantia nigra by integrating scRNA-seq datasets.

We also linked single-cell epigenomic and gene expression states by integrating transcriptomic and DNA methylation data from the mouse frontal cortex using LIGER. This joint analysis aided in the interpretation of populations difficult to identify from methylation alone and increased our sensitivity for detecting rare cell types. It further allowed us to investigate the epigenetic regulation of gene expression for each jointly defined cell type.

In addition, we jointly defined cell populations using scRNA-seq profiles and spatial transcriptomic data (StarMAP) from mouse frontal cortex. This joint analysis not only enabled inference of spatial information for gene expression profiles from dissociated cells, but also increased the sensitivity for identifying cell clusters from the *in situ* data.

### Applications of the method

Tran and Shekhar et al. recently used LIGER in their study of neuronal type-specific response to injury^3^. They focused on the adult mouse retinal ganglion cells (RGCs) and investigated the resilience of RGC types following optic nerve crush (ONC), a common model of traumatic axonal injury. The authors employed scRNA-seq to profile the injured RGCs at different time points post ONC. They used LIGER to develop a common taxonomy of cell types that was robust to time of injury, mouse strain, and batch effects. The capability of LIGER to distinguish shared features such as RGC type-specific expression pattern and dataset-specific features--such as injury-related changes--along the time course enabled discovery of RGC type-specific molecular signatures related to cell resilience and susceptibility to injury. More recently, Krienen et al. applied LIGER to probe interneuron cell types and their gene expression patterns across multiple species, including humans, macaques, marmosets, and mice^4^. The authors used LIGER to jointly define interneuron cell types across species and brain regions from Drop-Seq data. The resulting joint analysis revealed shared cell types across species; an interneuron cell type that appears in humans and monkeys but not mice; and species-specific gene expression differences within shared cell types.

### Comparison with other methods

There are several existing methods for single-cell data integration, including Scanorama, Seurat, Conos, and Harmony. Scanorama is a method designed for scRNA-seq dataset integration and batch correction and was introduced by Hie et al. in 2019^5^. Inspired by the idea of stitching images into a panoramic picture, the key strategy of this method is to identify the common cell types shared in all pairs of dataset by finding the mutual nearest neighbors in a low-dimensional space obtained by singular value decomposition. Then pairs of datasets are merged into a “cellular panorama” based on their matched cells. The panoramic stitching approach requires that each of the input datasets shares at least one cell type with at least one of the others.

The Kharchenko group developed a three-phase approach for clustering on networks of samples (Conos) from multiple scRNA-seq datasets^6^. The input datasets are filtered and normalized in Conos phase I. In phase II, pairwise comparison of the datasets using PCA embeddings is performed to obtain the initial mapping between the cell samples. In the last phase of Conos, a joint graph is constructed based on the inter-sample and intra-sample edges. The joint graph can be used for downstream analysis such as community detection, visualization, or label and expression value propagation.

Korsunsky and colleagues developed a multi-dataset integration algorithm, Harmony, in their recent publication^7^. Harmony takes an initial PCA embedding of scRNA-seq expression matrices as input and learns a batch-corrected embedding that allows for downstream analysis, including visualization and clustering. This is accomplished by an iterative process of soft clustering, calculating the centroids of the resulting clusters, computing cluster-specific correction vectors based on the centroids, and correcting the cells using the obtained correction vectors. The process is repeated until the devised objective, maximizing both the data integration and separation of cell types, is reached.

Scanorama, Harmony, and Conos are designed primarily for integrating multiple RNA datasets, but Seurat^8^ is the most similar method to LIGER in that it is intended for integrating multiple data modalities, including RNA, ATAC, and spatial transcriptomic data. However, there are some key differences. Seurat performs “label transfer” between reference and query datasets rather than joint cell type definition when integrating RNA and ATAC or spatial transcriptomic and dissociated data. Additionally, Seurat uses a completely shared latent space (CCA) to identify corresponding cells, rather than a space that includes dataset-specific differences. In contrast, LIGER’s dimensionality reduction strategy captures both shared signals and dataset-specific differences per cell type directly in the factorization. Additionally, the metagene factors learned by LIGER are often interpretable in terms of specific biological or technical sources of variation, whereas each of the principal components or canonical components calculated by Seurat often contains a mixture of signals. Finally, Seurat “corrects” the gene expression values using nearest neighbor relationships (“anchors”), whereas LIGER leaves the original expression data unchanged.

A recent paper performed systematic benchmarking of 14 single-cell data integration methods and recommended Harmony, LIGER, and Seurat as the top-performing and most robust approaches^9^. The paper benchmarked the 14 approaches on 10 different single-cell datasets using multiple metrics, including kBET^10^, average silhouette width, and adjusted rand index. Harmony, LIGER, and Seurat outperformed the other tested methods across a range of scenarios in their ability to align datasets while maintaining correct cell type distinctions.

### Experimental design

The design of single-cell sequencing experiments is complex, involving many biological and experimental considerations beyond the scope of this protocol. However, we will highlight two salient issues that affect the joint cell type analysis we describe in this protocol. First, whenever possible, biological and technical batches should not be confounded. For example, in a cross-species analysis of mouse and human brain cells, both mouse and human cells should be extracted using the same protocols; if whole-cell transcriptomes are extracted from the mouse cells, but only nuclear RNA is extracted from the human cells, the biological variable (species) is confounded by a technical variable (whole-cell versus nuclear extraction protocol). It may still be possible, using LIGER, to identify shared cell types and gene expression signatures using such a batch design, but disentangling biological from technical differences will be challenging. Similarly, if jointly defining cell types using scRNA-seq and snATAC-seq, use cells from the same tissue sample for both protocols. Otherwise, biological differences in cell composition or genotype may confound the joint analysis. For human studies comparing many individuals, genetic variation can be used to multiplex cells from multiple donors within a single batch, allowing reliable determination of cell-type-specific inter-individual variation without uncontrolled technical variables^11^.

A second consideration is the trade-off between the number of cells sequenced and sequencing depth per cell. Although the decision depends on the biological questions that the researcher wants to answer, it is generally preferable to sequence more cells rather than more reads per cell. A recent analysis paper found that, for droplet single-cell RNA-seq protocols, sequencing more than 15,000 reads per cell provides only small benefit^12^. Sequencing large numbers of cells is especially important for characterizing rare cell types, such as the recently discovered pulmonary ionocyte^13,14^. However, some biological applications, such as studying alternative splicing^15^ or characterizing important low-expression genes such as G-protein coupled receptors and transcription factors, will require a high depth of coverage per cell. Furthermore, the choice of library preparation protocol may also influence the decision about cells vs. reads, because protocols (such as SMART-seq) that sample from all positions within a transcript benefit more from increased coverage than poly(A) priming protocols that capture only the ends of transcripts.

### Limitations

LIGER should not be used when there is no biological similarity expected between datasets. For example, there is nothing to be gained by trying to integrate single-cell data from HeLA cells with data from HEK293 cells. Doing so will not yield significant biological insights and may result in false positive results, although LIGER is designed to minimize such spurious alignment. Additionally, LIGER relies on corresponding features shared across all datasets to be analyzed. Thus, it cannot be directly applied to integrate single-cell gene expression data with single-cell morphological data, chromatin conformation data, or metabolomic data, whose features have no clearly defined relationship with the expression of individual genes.

### Materials

- Hardware: a personal computer with internet connection
- Software:
  ∘ RStudio or R command line (version 3.5 or greater)
  ∘ bedmap (if calculating gene body counts from raw snATAC-seq data)
- Input data:
  ∘ Two or more matrices, each containing gene-level counts across a set of single cells
  ∘ A common starting point is the output from the 10X Genomics Cellranger pipeline, which outputs a gene expression matrix in matrix market (.mtx) format.

## Procedure

### Joint definition of cell types from multiple scRNA-seq datasets

This protocol demonstrates the R commands to run the LIGER package on a sample dataset consisting of two single-cell RNA-seq experiments. These commands can be run from the R command line interface or the RStudio integrated development environment.

#### Stage I: Preprocessing and Normalization (3 - 5 seconds)

**1**. For the first portion of this protocol, we will be integrating published data^11^ from control and interferon-stimulated peripheral blood mononuclear cells (PBMC). This dataset was originally in the form of output from the 10X Cellranger pipeline. We directly load downsampled versions of the control and stimulated DGEs here.

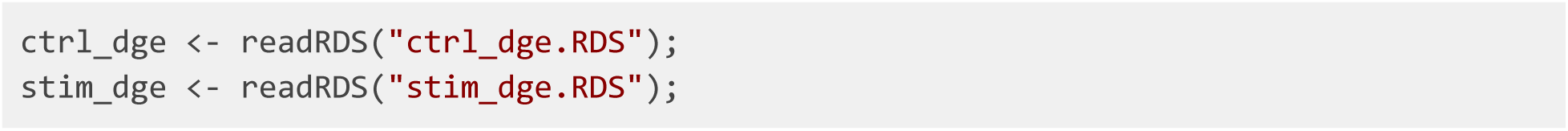

For 10X CellRanger output, we can instead use the ‘read10X’ function, which generates a matrix or list of matrices directly from the output directories.

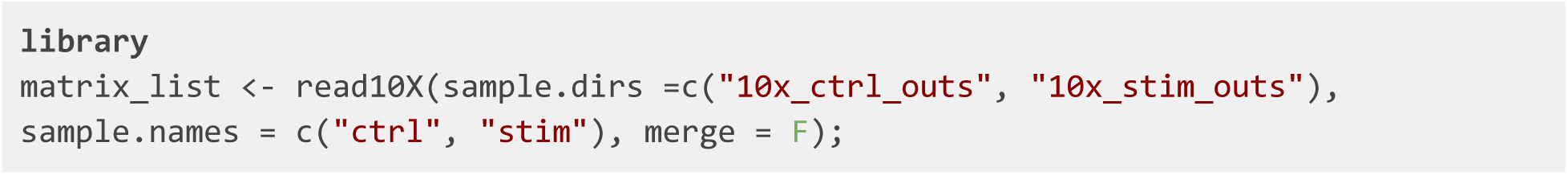

**?TROUBLESHOOTING**

**2**. With the digital gene expression matrices for both datasets, we can initialize a LIGER object using the *createLiger* function.

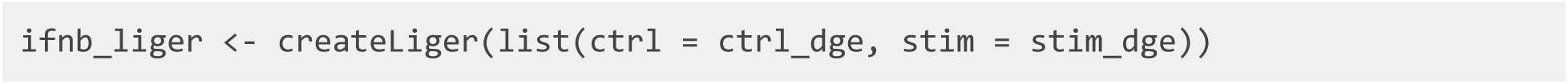

*ifnb_liger* now contains two datasets in its *raw.data* slot, ctrl and stim. We can run the rest of the analysis on this LIGER object.

**?TROUBLESHOOTING**

**3**. Before we can perform matrix factorization to integrate our datasets, we must run several preprocessing steps to normalize expression data to account for differences in sequencing depth and efficiency between cells, identify variably expressed genes, and scale the data so that each gene has the same variance. Note that because nonnegative matrix factorization requires positive values, we do not center the data by subtracting the mean. We also do not log transform the data.

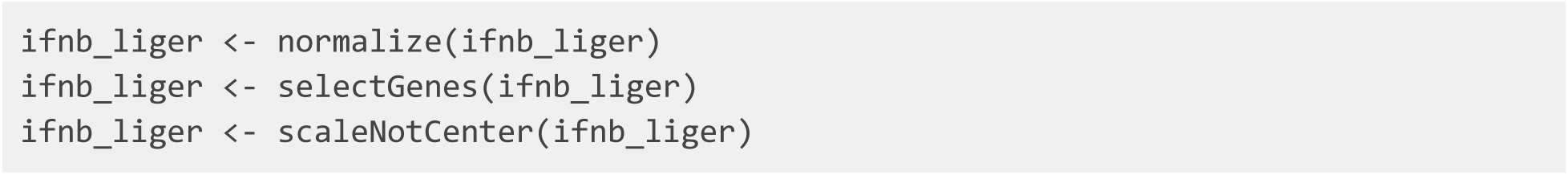

**?TROUBLESHOOTING**

#### Stage II: Joint Matrix Factorization (3-10 minutes)

**4**. We are now able to run integrative non-negative matrix factorization on the normalized and scaled datasets. The key parameter for this analysis is *k*, the number of matrix factors (analogous to the number of principal components in PCA). In general, we find that a value of *k* between 20 and 40 is suitable for most analyses and that results are robust for choice of *k*. Because LIGER is an unsupervised, exploratory approach, there is no single “right” value for *k*, and in practice, users choose *k* from a combination of biological prior knowledge and other information.

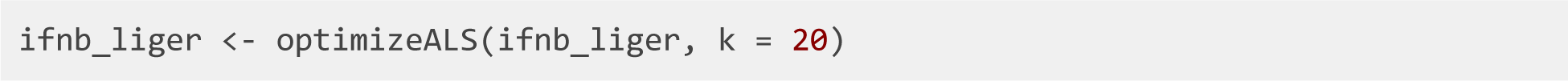

The optimization yields several low-dimensional matrices, including the *H* matrix of metagene expression levels for each cell, the *W* matrix of shared metagenes, and the *V* matrices of dataset-specific metagenes.

Please note that the time required of this step is highly dependent on the size of the datasets being used. For datasets of about 30,000 cells or less, this step should take less than 30 minutes.

#### Stage III: Quantile Normalization and Joint Clustering (1-2 minutes)

**5**. We can now use the resulting factors to jointly cluster cells and perform quantile normalization by dataset, factor, and cluster to fully integrate the datasets. All of this functionality is encapsulated within the *quantile_norm* function, which uses maximum factor assignment followed by refinement using a k-nearest neighbors graph.

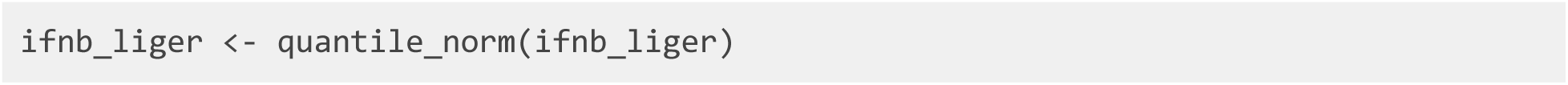

**6**. The *quantile_norm* procedure produces joint clustering assignments and a low-dimensional representation that integrates the datasets together. These joint clusters directly from iNMF can be used for downstream analyses (see below). Alternatively, we can also run Louvain community detection, an algorithm commonly used for single-cell data, on the normalized cell factors. The Louvain algorithm excels at merging small clusters into broad cell classes and thus may be more desirable in some cases than the maximum factor assignments produced directly by iNMF.

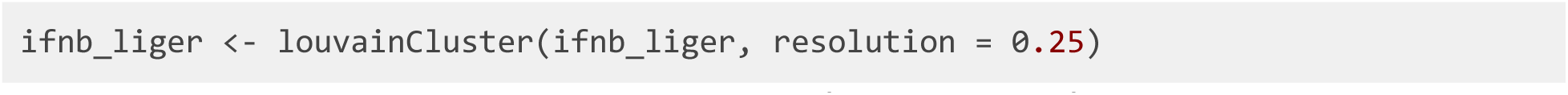

#### Stage IV: Visualization and Downstream Analysis (25-40 seconds)

**7**. To visualize the clustering of cells graphically, we can project the normalized cell factors to two or three dimensions. LIGER supports both t-SNE and UMAP for this purpose. Note that if both techniques are run, the object will only hold the results from the most recent.

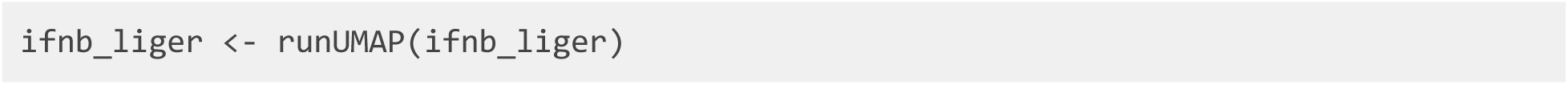

The LIGER package implements a variety of utilities for visualization and analysis of clustering, gene expression across datasets, and comparisons of cluster assignments. We will summarize several here.

**8**. *plotByDatasetAndCluster* plots two images using the coordinates generated by t-SNE or UMAP in the previous step. The first plot colors cells by dataset of origin, and the second colors cells by joint cluster assignment. The plots provide visual confirmation that the datasets are well aligned and the clusters are consistent with the structure of the data as revealed by UMAP (**Figure 2**).

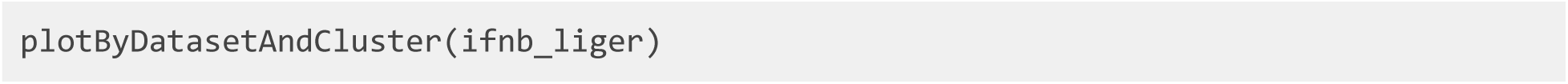

**Figure 1:**
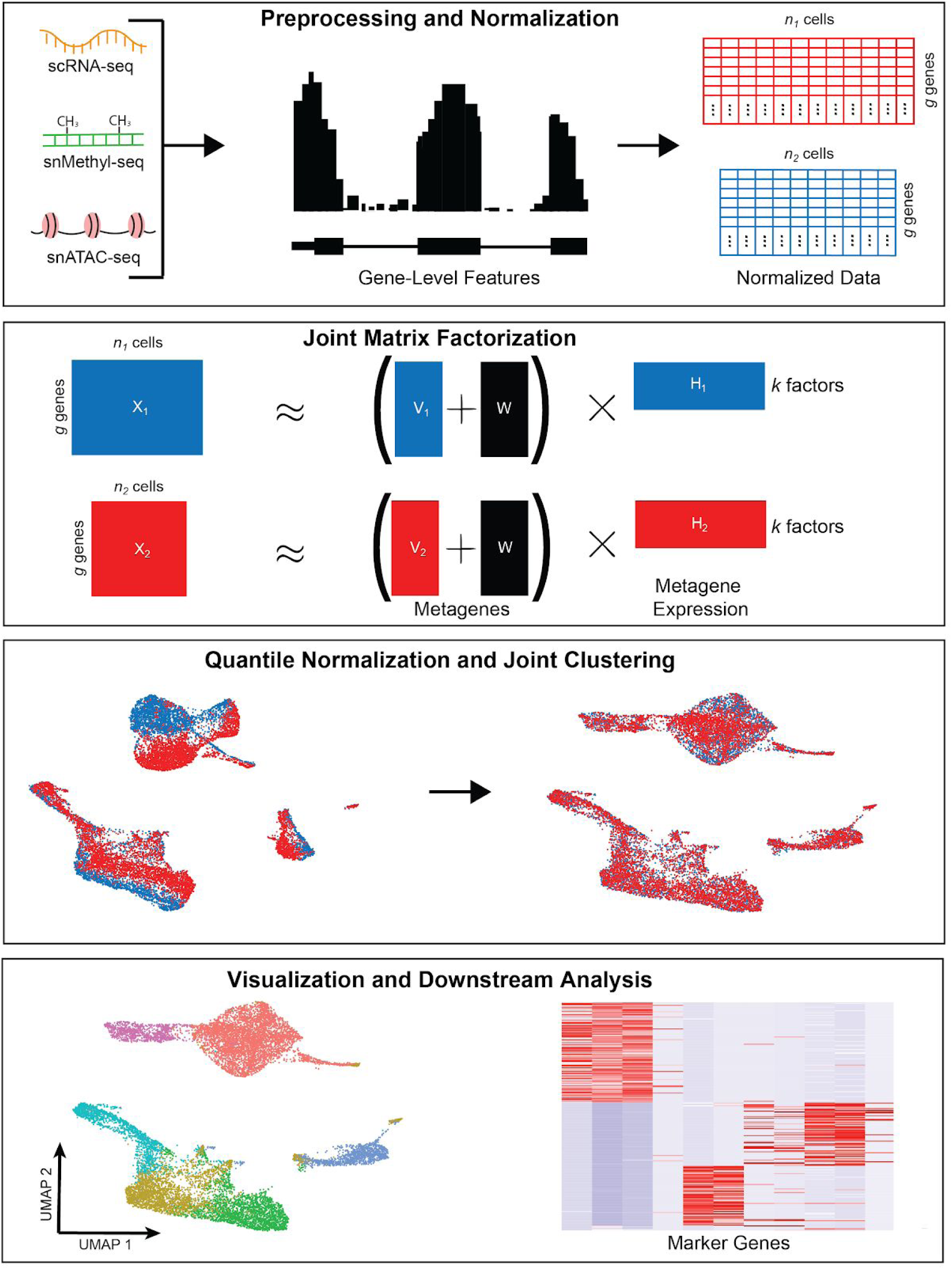
Diagram of high-level protocol steps.

**Figure 2:**
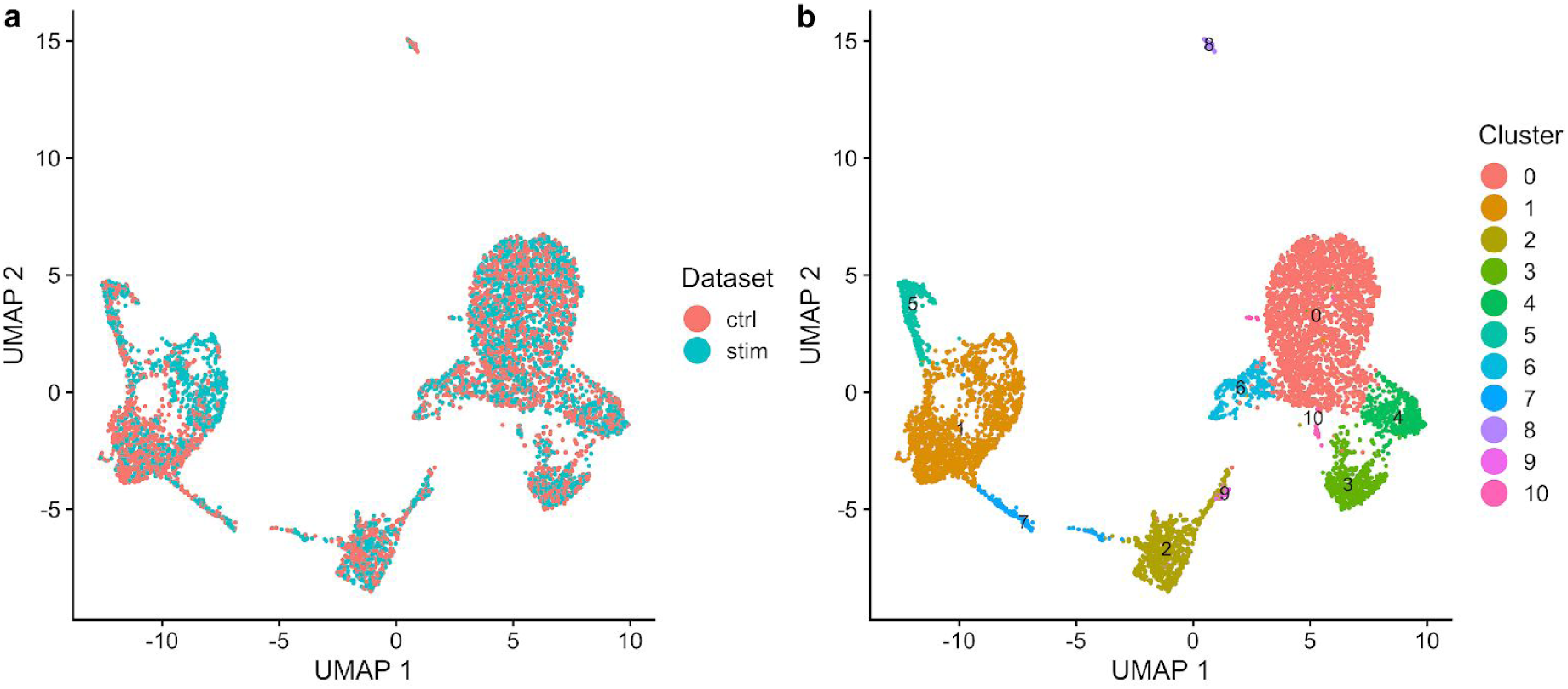
LIGER jointly identifies clusters from across single-cell RNA-seq datasets. **a,b**, Uniform manifold approximation and projection (UMAP) plots of a LIGER analysis of 3000 control and 3000 interferon-beta stimulated PBMCs profiled by scRNA-seq, colored by dataset (**a**) and LIGER joint cluster assignment (**b**).

To directly study the impact of factors on the clustering and determine what genes load most highly on each factor, we use the *plotGeneLoadings* function, which returns plots of metagene expression levels on the dimensionally reduced graphs and gene loading values by dataset for each metagene (**Figure 3**).

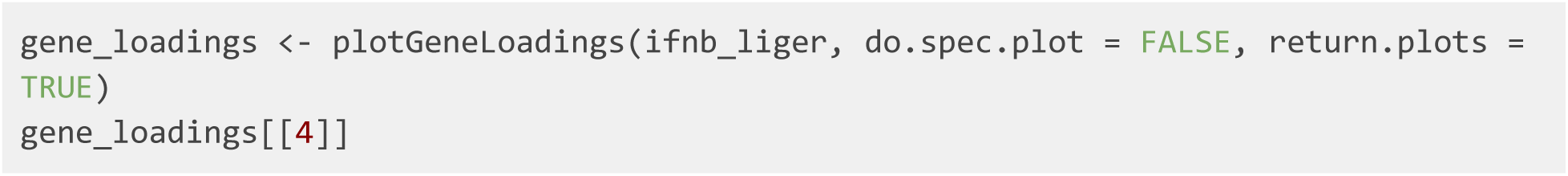

**Figure 3:**
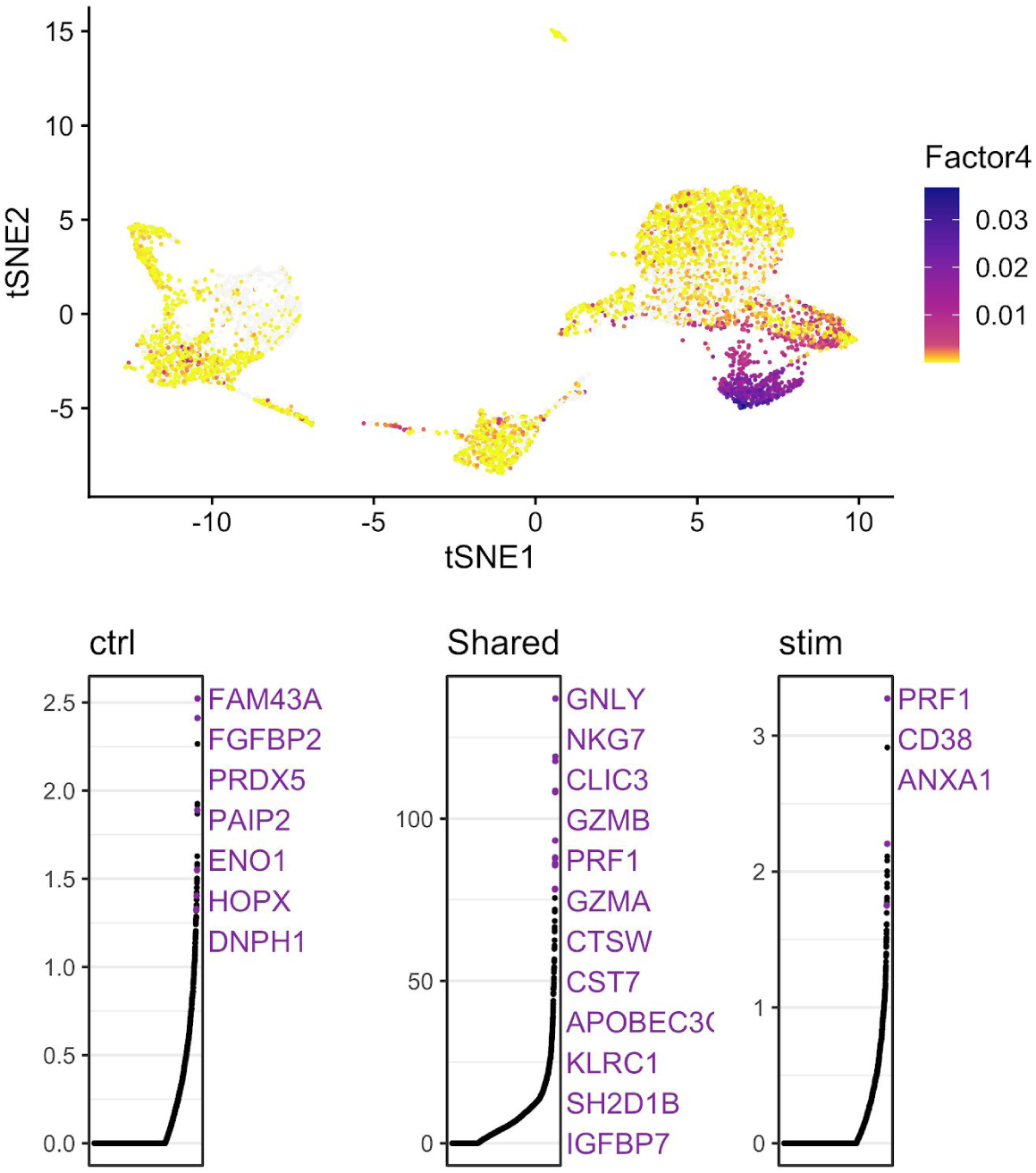
LIGER enables metagene and dataset specific analysis of PBMC data. UMAP plots showing metagene expression levels (factor loading values for each cell; top) and gene loadings (on a particular metagene; bottom) for Factor 4, which specifically loads on Cluster 3. In gene loading plots, gene names are sorted in decreasing order of magnitude of their factor loading contribution and correspond to colored points in scatterplots. Plots are organized to show the dataset-specific metagene values for control cells, the shared metagene values common to all datasets and the dataset-specific metagene values for interferon-stimulated cells.

Using the *runWilcoxon* function, we can next identify gene markers for all clusters. We can also compare expression within each cluster across datasets, which in this case reveals markers of interferon-beta stimulation. The function returns a table of data that allows us to determine the significance of each gene’s differential expression, including log fold change, area under the curve and *p*-value (**Table 1**).

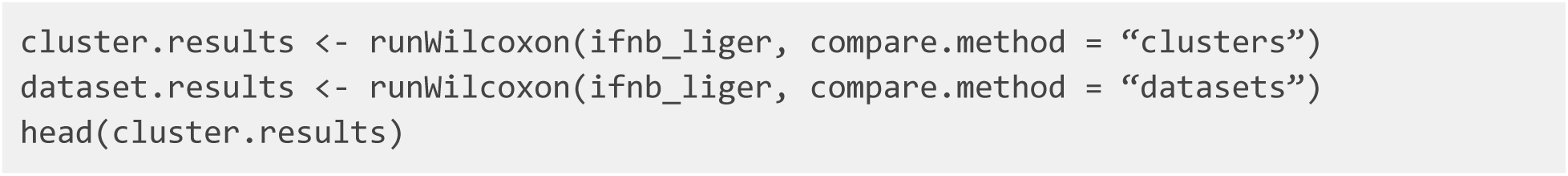

**Table 1:**
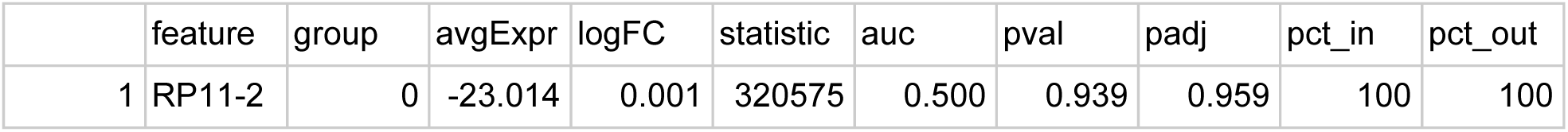
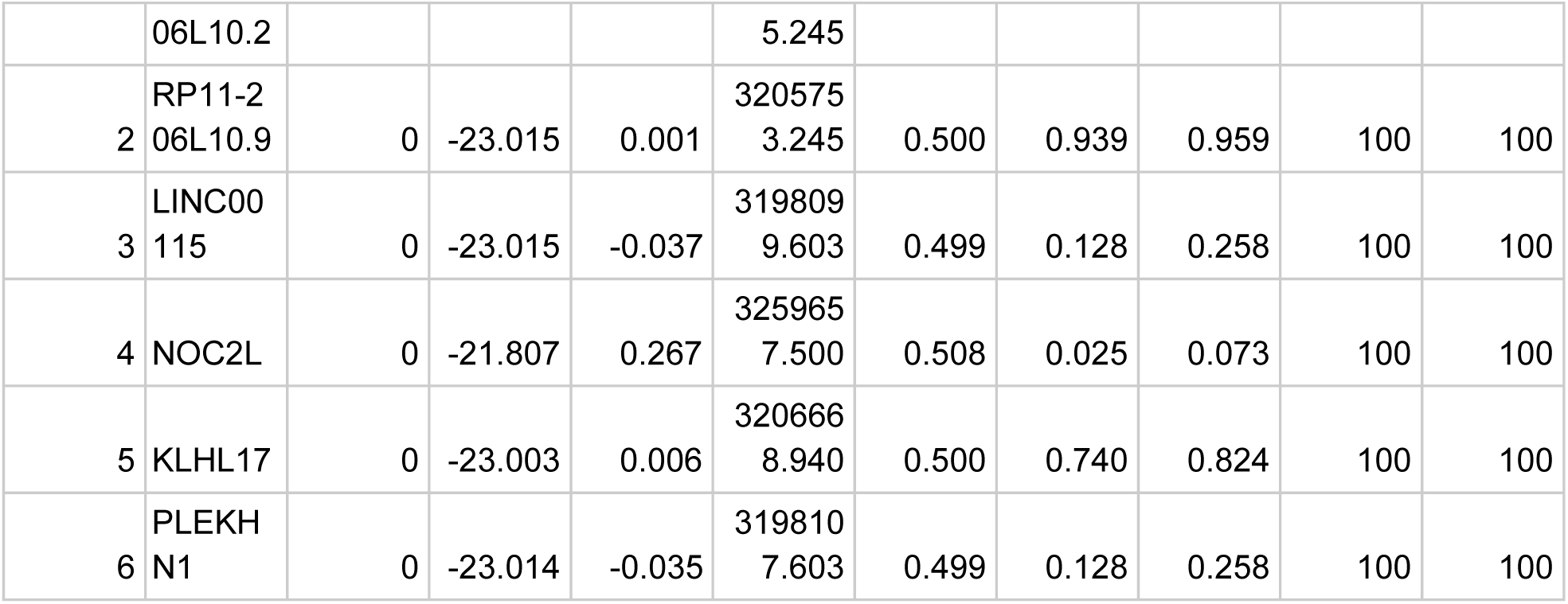
Wilcoxon test results indicating shared cluster markers across datasets.

The number of marker genes identified by *runWilcoxon* varies and depends on the datasets used. The function outputs a data frame that the user can then filter to select markers that are statistically and biologically significant. For example, one strategy is to filter the output by taking markers which have *padj* (Benjamini-Hochberg adjusted *p*-value) less than 0.05 and *logFC* (log fold change between observations in-group versus out-group) larger than 3:

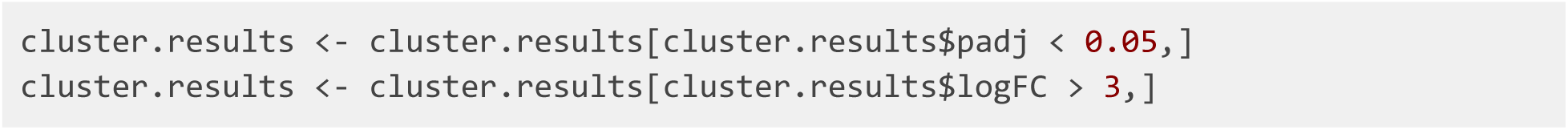

We can then re-sort the markers by *padj* value in ascending order and choose the top 100 for each cell type. For example, we can subset and re-sort the output for *Cluster 3* and take the top 20 markers by typing these commands (**Table 2**):

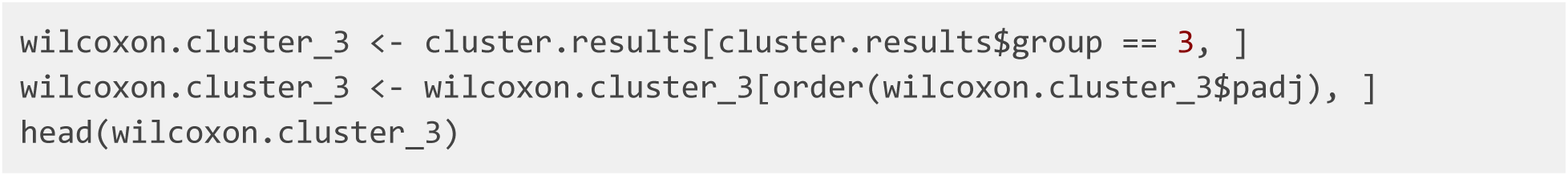

**Table 2:** Top shared cluster markers from the Wilcoxon test on IFNB dataset.

|  | feature | group | avgExpr | logFC | statistic | auc | pval | padj | pct_in | pct_out |
| --- | --- | --- | --- | --- | --- | --- | --- | --- | --- | --- |
| 41861 | GNLY | 3 | -5.282466 | 16.1473<br>23 | 2379942 | 0.96476<br>42 | 0 | 0 | 100 | 100 |
| 46904 | CLIC3 | 3 | -12.96207<br>0 | 9.48559<br>9 | 1946458 | 0.78904<br>17 | 0 | 0 | 100 | 100 |
| 47130 | PRF1 | 3 | -12.26057<br>3 | 9.83008<br>2 | 1970124 | 0.79863<br>52 | 0 | 0 | 100 | 100 |
| 49239 | GZMB | 3 | -7.840488 | 13.6971<br>78 | 2231966 | 0.90477<br>89 | 0 | 0 | 100 | 100 |
| 52832 | NKG7 | 3 | -6.594620 | 14.4331<br>85 | 2324832 | 0.94242<br>43 | 0 | 0 | 100 | 100 |
| 48310 | KLRC1 | 3 | -18.00005<br>9 | 4.91623<br>9 | 1606731 | 0.65132<br>52 | 1.83940<br>5e-289 | 4.09911<br>4e-286 | 100 | 100 |

We can visualize the expression profiles of individual genes, such as the differentially expressed genes that we just identified. This allows us to visually confirm the cluster- or dataset-specific expression patterns of marker genes. The UMAP plots of *PRF1* expression indicate that this gene is a specific marker for Cluster 3, with high values in this cell group and low values elsewhere (**Figure 4**). Meanwhile, we can also see that the distributions are very similar between the control and interferon-stimulated datasets, indicating that LIGER has properly aligned these two datasets.

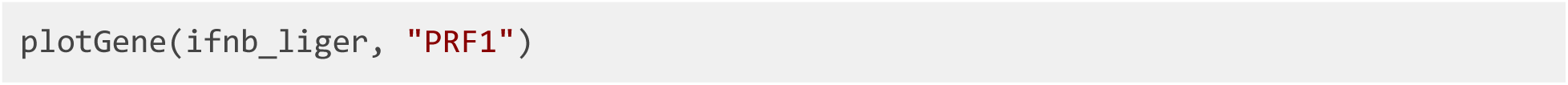

**Figure 4:**
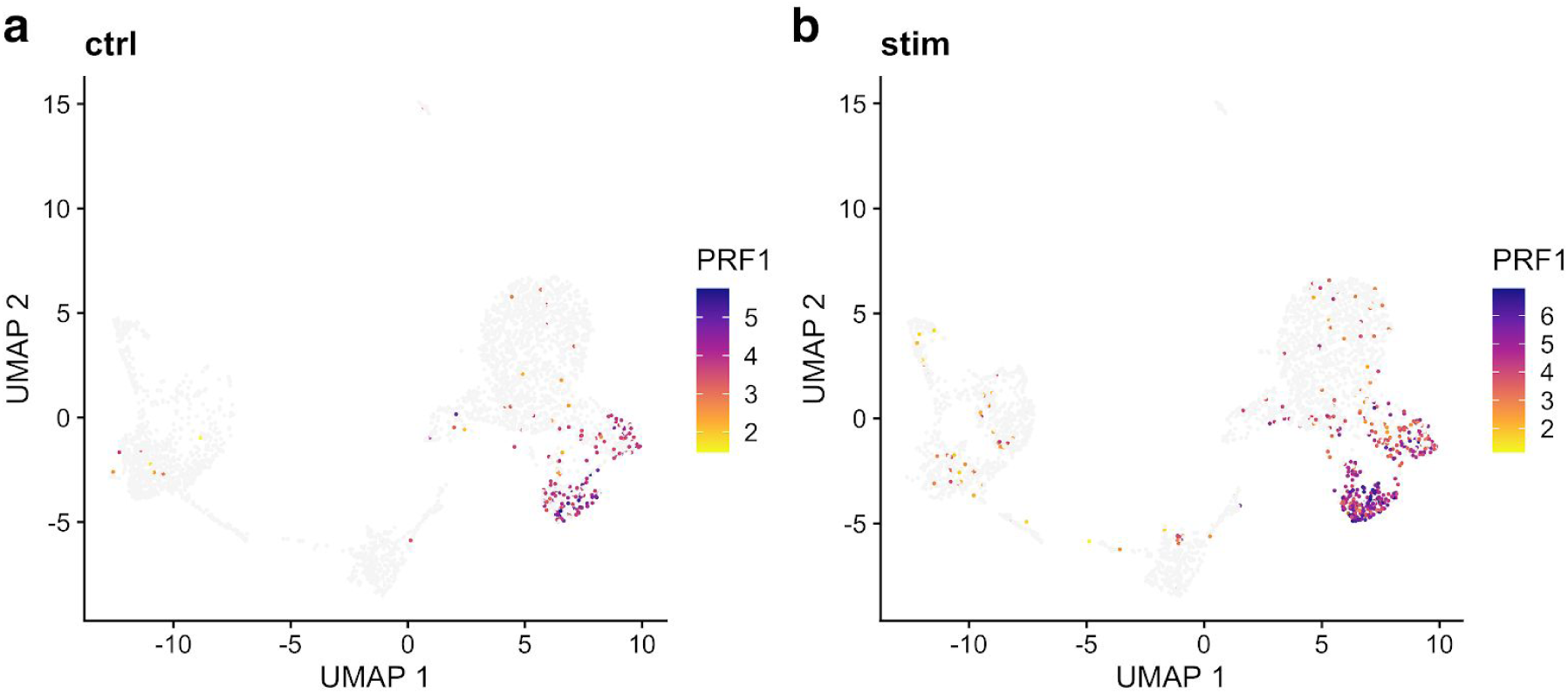
Marker gene identified by LIGER shows consistent cell-type-specific expression across datasets. **a,b**, UMAP representations of expression for gene *PRF1*, a marker gene of cluster 3, in control (**a**) and interferon-beta stimulated (**b**) PBMCs exhibit similar distributions.

We can also use *plotGene* to inspect genes with expression that differs within a cluster across datasets.

### Joint definition of cell types from single-cell gene expression and single-nucleus chromatin accessibility data (Human bone marrow mononuclear cells)

In this section, we will demonstrate LIGER’s ability to jointly define cell types by leveraging multiple single-cell modalities. We integrate published human bone marrow mononuclear cell (BMMC) data ^16^ profiled by single-cell RNA-seq and single-nucleus ATAC-seq to enable cell type definitions that incorporate both gene expression and chromatin accessibility data. Such joint analysis allows not only the taxonomic categorization of cell types, but also a deeper understanding of their underlying regulatory networks. The pipeline for jointly analyzing scRNA-seq and snATAC-seq is similar to that for integrating multiple scRNA-seq datasets in that both rely on joint matrix factorization and quantile normalization. The main differences are: (1) snATAC-seq data needs to be processed into gene-level values; (2) gene selection is performed on the scRNA-seq data only; and (3) downstream analyses can use both gene-level and intergenic information.

#### Stage I: Preprocessing and Normalization (40 - 50 minutes)

In order to jointly analyze scRNA and snATAC-seq data, we first need to transform the snATAC-seq data--a genome-wide epigenomic measurement--into gene-level counts which are comparable to gene expression data from snRNA-seq. Most previous single-cell studies have used an approach inspired by traditional bulk ATAC-seq analysis: identifying chromatin accessibility peaks, then summing together all peaks that overlap each gene. This strategy is also appealing because the 10X CellRanger pipeline, a commonly used commercial package, automatically outputs such peak counts. However, we find this peak summing strategy undesirable because: (1) peak calling is performed using all cells, which biases against rare cell populations; (2) gene body accessibility is often more diffuse than that of specific regulatory elements, and thus may be missed by peak calling algorithms; and (3) information from reads outside of peaks is discarded, further reducing the amount of data in the already sparse measurements. Instead of summing peak counts, we find that the simplest possible strategy seems to work well: counting the total number of ATAC-seq reads within the gene body and promoter region (typically 3 kb upstream) of each gene in each cell.

Note that in this part, we included the details of running this preprocessing workflow for only one sample. Users should re-run this counting step multiple times for more than one snATAC-seq sample.

Note also that several commands need to be run through the **Command Line Interface** instead of the R Console or IDE (RStudio). We also employ the *bedmap* command from the BEDOPS tool to make a list of cell barcodes that overlap each gene and promoter. The gene body and promoter indexes are .*bed* files, which indicate gene and promoter coordinates. Since *bedmap* expects sorted inputs, the fragment output from CellRanger, gene body and promoter indexes should all be sorted.

We show below how to perform these steps for snATAC-seq data generated by the 10X Chromium system, the most widely used snATAC-seq platform. The input for this process is the file *fragments.tsv* output by CellRanger, which contains all ATAC reads that passed filtering steps.

**1**. We must first sort *fragments.tsv* by chromosome, start, and end position using the *sort* command line utility. The *-k* option lets the user sort the file on a certain column; including multiple *-k* options allows sorting by multiple columns simultaneously. The *n* behind *-k* stands for ‘numeric ordering’. Here the sorted .*bed* file order is defined first by lexicographic chromosome order (using the parameter *-k1,1*), then by ascending integer start coordinate order (using parameter *-k2,2n*), and finally by ascending integer end coordinate order (using parameter *-k3,3n*). Note that this step may take a while, since the input fragment file is usually very large (for example, a typical fragment file of 4-5 GB can take about 40 minutes).

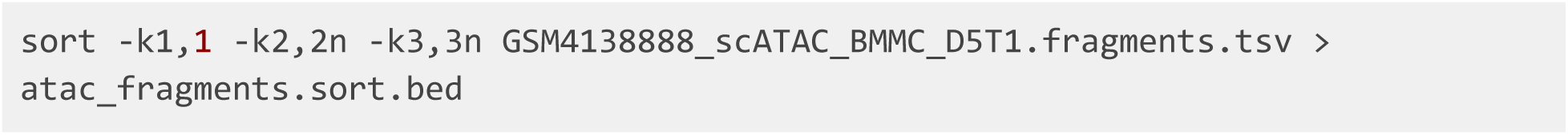

Gene body and promoter locations should also be sorted using the same strategy for sorting fragment output files:

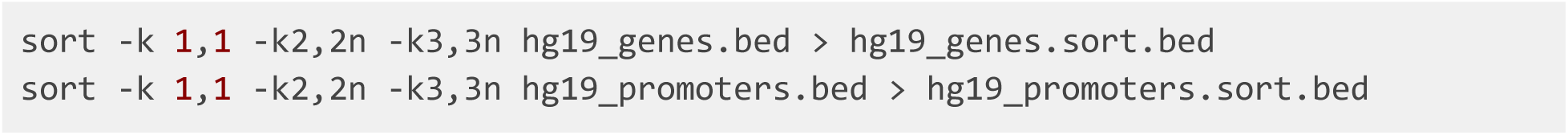

**2**. Use the *bedmap* command to determine which fragments overlap each gene body and promoter:

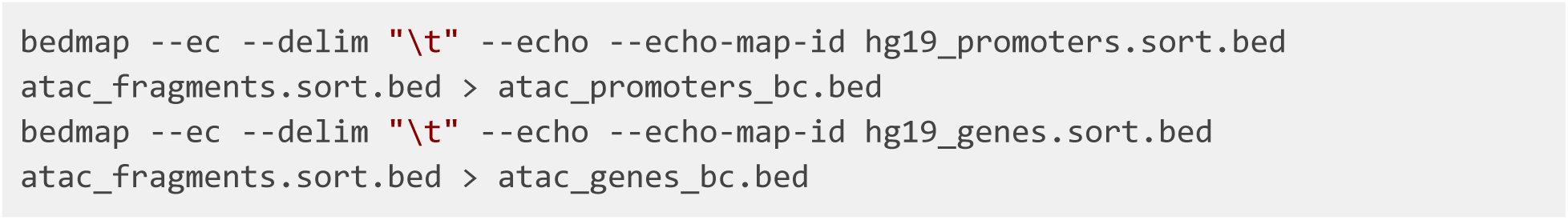

Important flags are as follows:

- *--delim*. This changes output delimiter from ‘|’ to specified delimiter between columns, which in our case is “\t”.
- *--ec*. Adding this will check input files to make sure that they are properly formatted and sorted.
- *--echo*. Adding this will print each line from reference file in output. The reference file in our case is gene or promoter index.
- *--echo-map-id*. Adding this will list IDs of all overlapping elements from mapping files, which in our case are cell barcodes from fragment files.

**3**. We then import the *bedmap* outputs into the R Console or RStudio. Note that the *as.is* option in *read.table* is specified to prevent the conversion of character columns to factor columns:

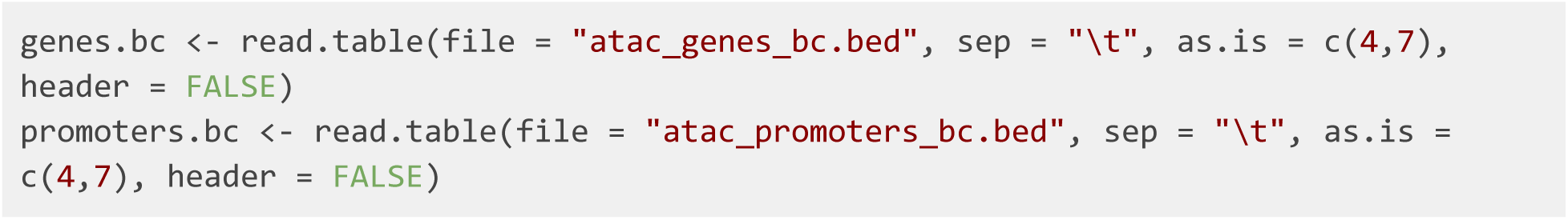

Cell barcodes are then split and extracted from the outputs. We recommend filtering barcodes that have a total number of reads lower than a certain threshold, for example, 1500. This threshold can be adjusted according to the size and quality of the samples.

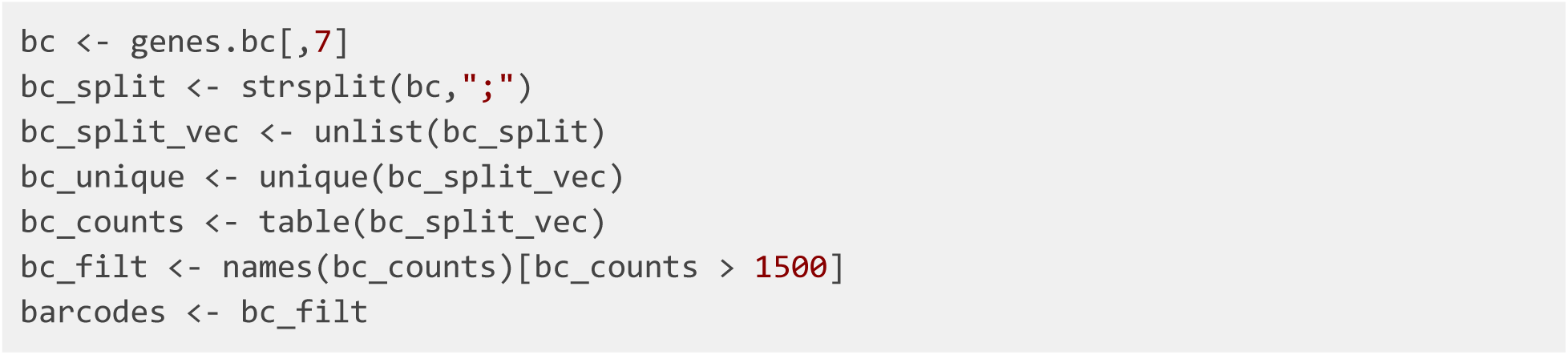

**4**. We can then use LIGER’s *makeFeatureMatrix* function to calculate accessibility counts for gene body and promoter individually. This function takes the output from *bedmap* and efficiently counts the number of fragments overlapping each gene and promoter. We could count the genes and promoters in a single step, but choose to calculate them separately in case it is necessary to look at gene or promoter accessibility individually in downstream analyses.

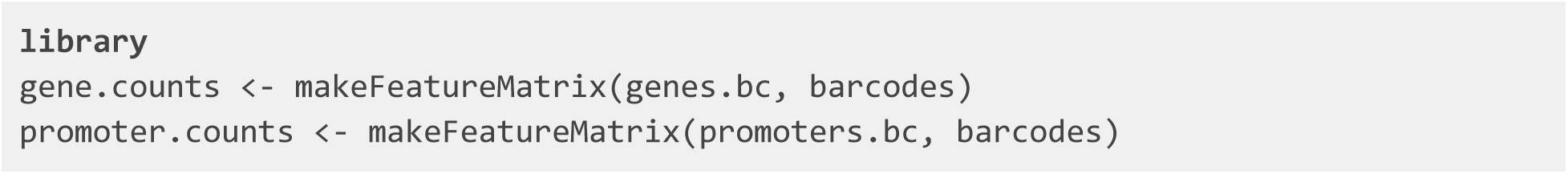

Next, these two count matrices need to be re-sorted by gene symbol. We then add the matrices together, yielding a single matrix of gene accessibility counts in each cell. Note that we also append the sample name to each cell barcode to avoid duplicate cell names across experiments.

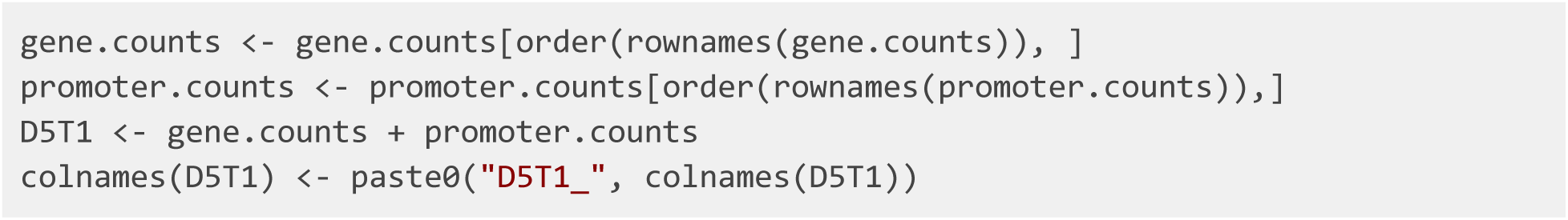

**?TROUBLESHOOTING**

**5**. Once the gene-level snATAC-seq counts are generated, the *read10X* function from LIGER can be used to read scRNA-seq count matrices output by CellRanger. You can pass in a directory (or a list of directories) containing raw outputs (for example, “*/Sample_1/outs/filtered_feature_bc_matrix*”) to the parameter *sample.dirs*. Next, a vector of names to use for the sample (or samples, corresponding to *sample.dirs*) should be passed to parameter *sample.names* as well. LIGER can also use data from any other protocol, as long as it is provided in a genes x cells R matrix format.

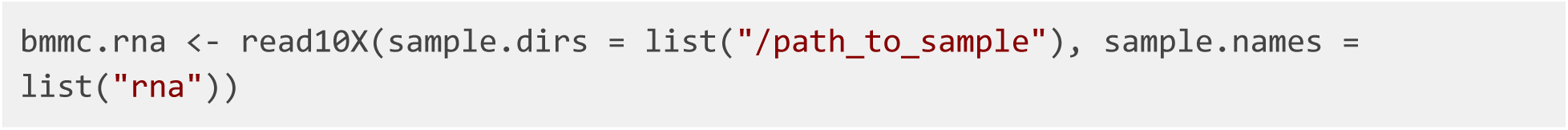

**6**. We can now create a LIGER object with the *createLiger* function. We also remove unneeded variables to conserve memory.

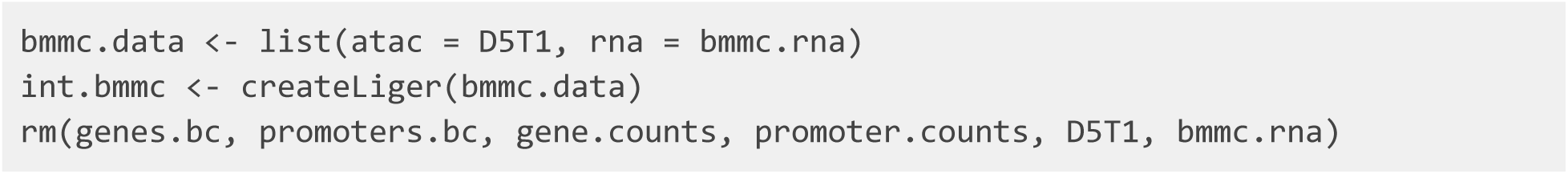

**?TROUBLESHOOTING**

**7**. Preprocessing steps are needed before running matrix factorization. Each dataset is normalized to account for differences in total gene-level counts across cells using the *normalize* function. Next, highly variable genes from each dataset are identified and combined for use in downstream analysis. Note that by setting the parameter *datasets.use* to 2, genes will be selected only from the scRNA-seq dataset (the second dataset) by the *selectGenes* function. We recommend not using the ATAC-seq data for variable gene selection because the statistical properties of the ATAC-seq data are very different from scRNA-seq, violating the assumptions made by the statistical model we developed for selecting genes from RNA data. Finally, the *scaleNotCenter* function scales normalized datasets without centering by the mean, giving the nonnegative input data required by iNMF.

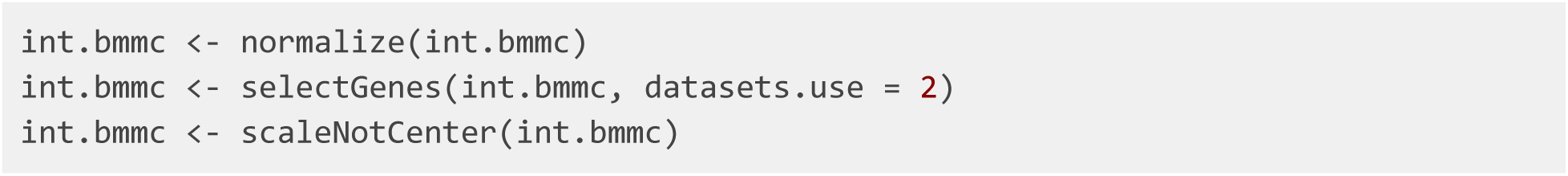

**?TROUBLESHOOTING**

#### Stage II: Joint Matrix Factorization (3 - 10 minutes)

**8**. We next perform joint matrix factorization (iNMF) on the normalized and scaled RNA and ATAC data. This step calculates metagenes--sets of co-expressed genes that distinguish cell populations--containing both shared and dataset-specific signals. The cells are then represented in terms of the “expression level” of each metagene, providing a low-dimensional representation that can be used for joint clustering and visualization. To run iNMF on the scaled datasets, we use the *optimizeALS* function with proper hyperparameter settings.

The important parameters are as follows:

- *k*. Integer value specifying the inner dimension of factorization, or number of factors. Higher *k* is recommended for datasets with more substructure. We find that a value of *k* in the range 20 - 40 works well for most datasets. Because this is an unsupervised, exploratory analysis, there is no single “right” value for *k*, and in practice, users choose *k* from a combination of biological prior knowledge and other information.
- *lambda*. This is a regularization parameter. Larger values penalize dataset-specific effects more strongly, causing the datasets to be better aligned, but possibly at the cost of higher reconstruction error. The default value is 5. We recommend using this value for most analyses, but find that it can be lowered to 1 in cases where the dataset differences are expected to be relatively small, such as scRNA-seq data from the same tissue but different individuals.
- *thresh*. This sets the convergence threshold. Lower values cause the algorithm to run longer. The default is 1e-6.
- *max.iters*. This variable sets the maximum number of iterations to perform. The default value is 30.

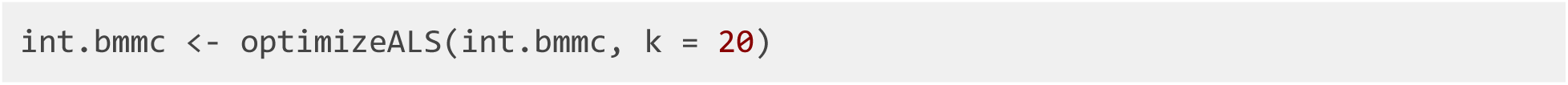

#### Stage III: Quantile Normalization and Joint Clustering (1 minute)

**9**. Using the metagene factors calculated by iNMF, we then assign each cell to the factor on which it has the highest loading, giving joint clusters that correspond across datasets. We then perform quantile normalization by dataset, factor, and cluster to fully integrate the datasets.

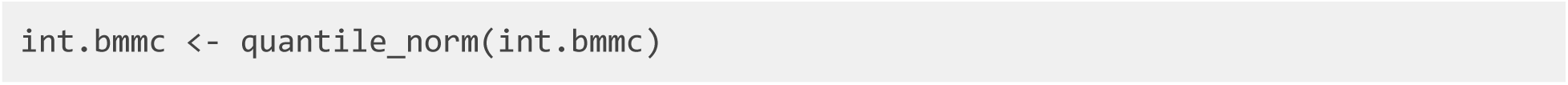

Important parameters of *quantile_norm* are as follows:

- *knn_k*. This sets the number of nearest neighbors for within-dataset KNN graph. The default is 20.
- *quantiles.* This sets the number of quantiles to use for quantile normalization. The default is 50.
- *min_cells*. This indicates the minimum number of cells to consider a cluster as shared across datasets. The default is 20.
- *dims.use*. This sets the indices of factors to use for quantile normalization. The user can pass in a vector of indices indicating specific factors. This is helpful for excluding factors capturing biological signals such as the cell cycle or technical signals such as mitochondrial genes. The default is all *k* of the factors.
- *do.center*. This indicates whether to center the data when scaling factors. The default is FALSE. This option should be set to TRUE when metagene loadings have a mean above zero, as with dense data such as DNA methylation.
- *max_sample*. This sets the maximum number of cells used for quantile normalization of each cluster and factor. The default is 1000.
- *refine.knn*. This indicates whether to increase robustness of cluster assignments using KNN graph. The default is TRUE.
- *eps*. This sets the error bound of the nearest neighbor search. The default is 0.9. Lower values give more accurate nearest neighbor graphs but take much longer to compute. We find that this parameter affects result quality very little.
- *ref_dataset*. This indicates the name of the dataset to be used as a reference for quantile normalization. By default, the dataset with the largest number of cells is used.

**10**. The *quantile_norm* function gives joint clusters that correspond across datasets, which are often completely satisfactory and sufficient for downstream analyses. However, if desired, after quantile normalization, users can additionally run the Louvain algorithm for community detection, which is widely used in single-cell analysis and excels at merging small clusters into broad cell classes. This can be achieved by running the *louvainCluster* function. Several tuning parameters, including *resolution, k*, and *prune* control the number of clusters produced by this function. For this dataset, we use a resolution of 0.2, which yields 18 clusters (see below).

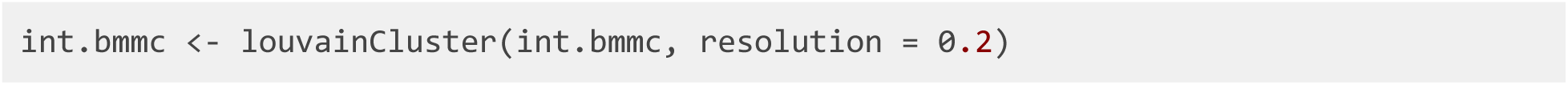

#### Stage IV: Visualization (2 - 3 minutes) and Downstream Analysis (30 - 40 minutes)

**11**. In order to visualize the clustering results, the user can use two dimensionality reduction methods supported by LIGER: t-SNE and UMAP. We find that often for datasets containing continuous variation such as cell differentiation, UMAP better preserves global relationships, whereas t-SNE works well for displaying discrete cell types, such as those in the brain. The UMAP algorithm (called by the *runUMAP* function) scales readily to large datasets. The *runTSNE* function also includes an option to use FFtSNE, a highly scalable implementation of t-SNE that can efficiently process large datasets. For the BMMC dataset, we expect to see continuous lineage transitions among the differentiating cells, so we use UMAP to visualize the data in two dimensions (**Figure 6**).

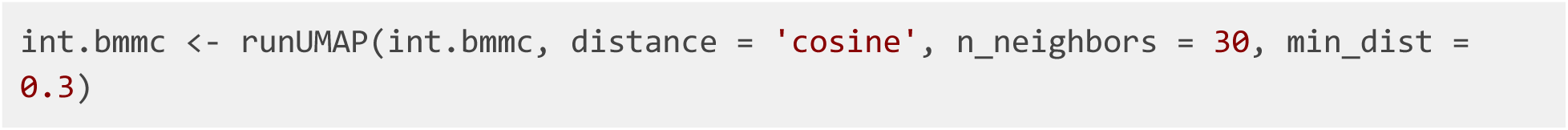

**Figure 5:**
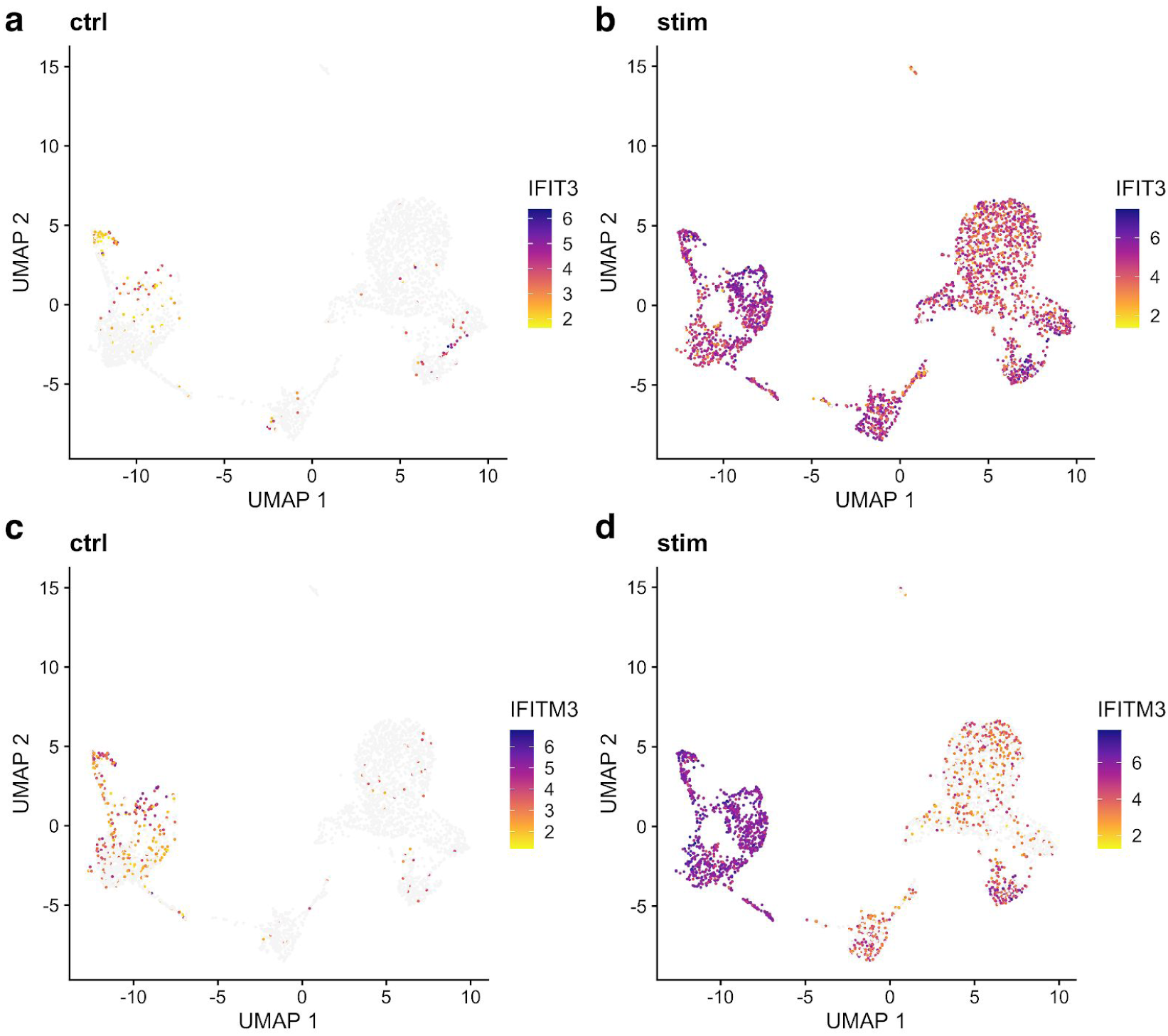
Marker genes identified by LIGER show expression differences across datasets. **a,b**, UMAP representations of expression for gene *IFIT3*, a marker gene of the interferon-stimulated dataset, shows low expression in control (**a**) and high expression in interferon-stimulated (**b**) PBMCs. **c,d**, UMAP representations of expression for gene *IFITM3*, a marker gene of cluster 1, in control (**c**) and IFNB-stimulated (**d**) PBMCs similarly shows more expression in interferon-stimulated cells.

**Figure 6:**
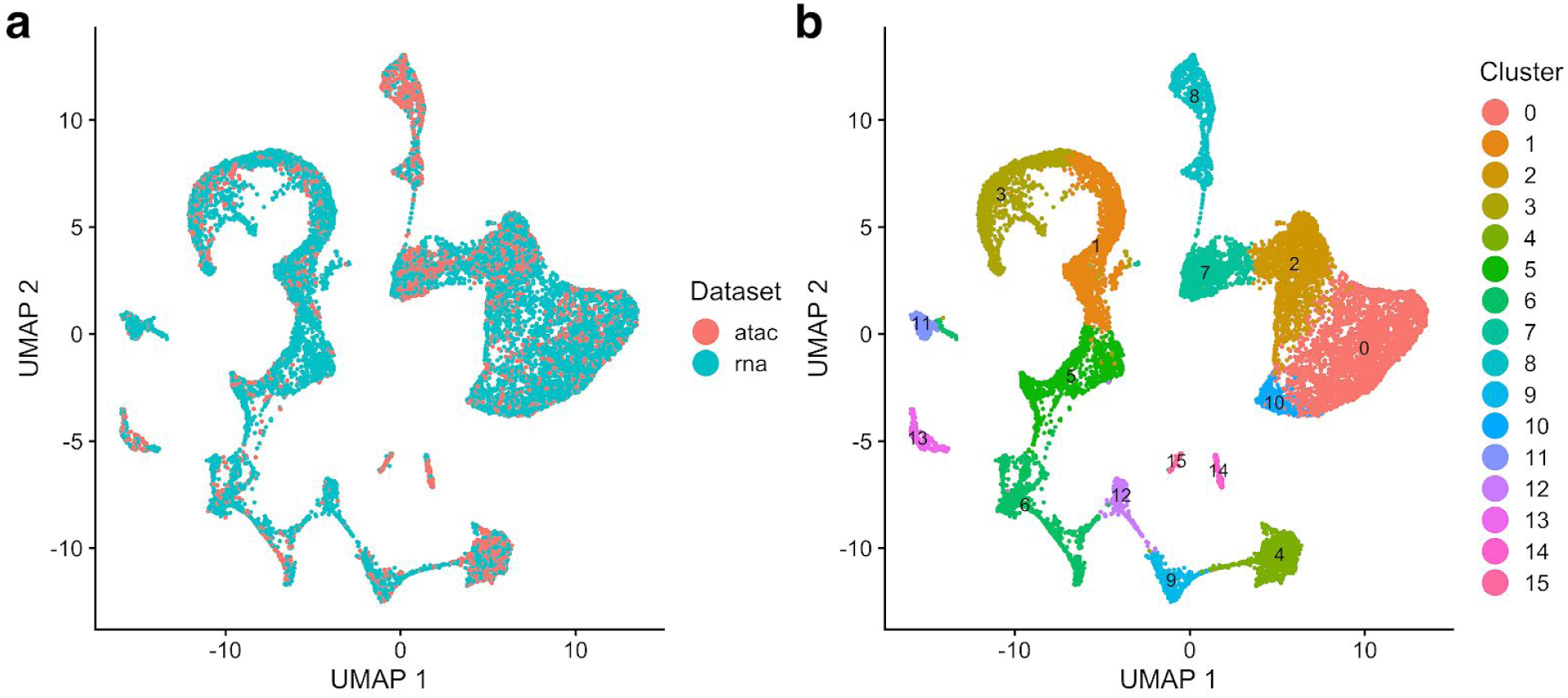
LIGER allows integrated alignment and clustering of BMMC data across technologies. **a,b**, UMAP plots of a LIGER analysis of 12,602 scRNA-seq profiles and 6,234 nuclei profiled by snATAC-seq, colored by technology (**a**) and LIGER cluster assignment (**b**).

**12**. We can then visualize each cell, colored by cluster or dataset.

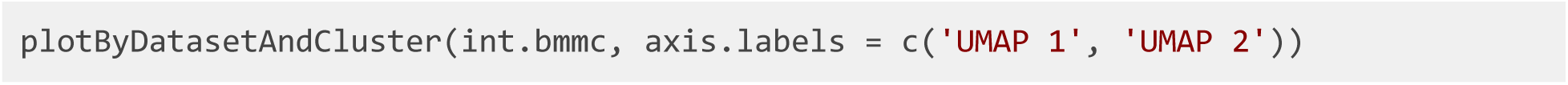

**13**. LIGER employs the Wilcoxon rank-sum test to identify marker genes that are differentially expressed in each cell type using the following settings. We provide parameters that allow the user to select which datasets to use (*data.use*) and whether to compare across clusters or across datasets within each cluster (*compare.method*). To identify marker genes for each cluster combining snATAC and scRNA profiles, typing in:

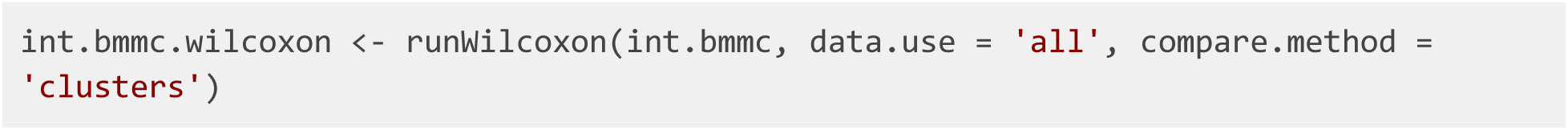

Important parameters of *runWilcoxon* are as follows:

- *data.use*. This selects which dataset (or set of datasets) to be included. The default is *‘all’* (using all the datasets).
- *compare.method*. This indicates whether to compare across clusters or across datasets within each cluster. Setting compare.method to *‘clusters’* compares each feature’s (genes, peaks, etc.) loading between clusters combining all datasets, which gives us the most specific features for each cluster. On the other hand, setting compare.method to *‘datasets’* gives us the features most differentially expressed across datasets for every cluster.

**14**. The number of marker genes identified by *runWilcoxon* varies and depends on the datasets used. The function outputs a data frame that the user can then filter to select markers that are statistically and biologically significant. For example, one strategy is to filter the output by taking markers which have *padj* (Benjamini-Hochberg adjusted *p*-value) less than 0.05 and *logFC* (log fold change between observations in group versus out) larger than 3:

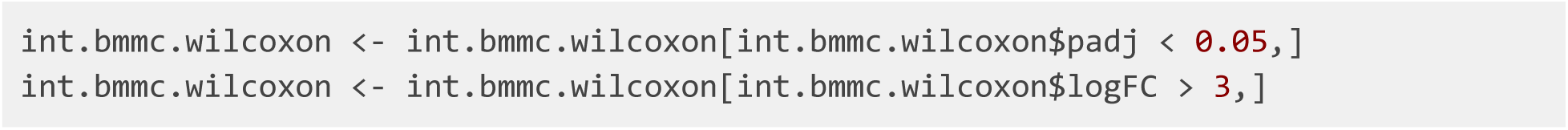

We can then sort the markers by *padj* value in ascending order and choose the top 100 for each cell type. For example, we can subset and re-sort the output for *Cluster 1* and take the top 20 markers by typing these commands:

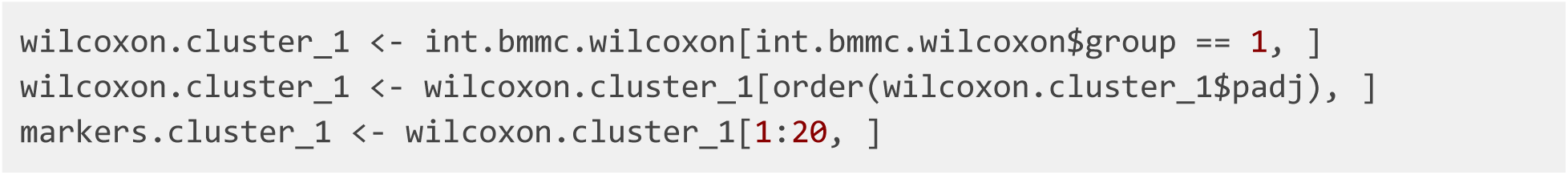

**15**. We also provide functions to check these markers by visualizing their expression across datasets. We can use the *plotGene* to visualize the expression or accessibility of a marker gene, which is helpful for visually confirming putative marker genes or investigating the distribution of known markers across the sequenced cells. Such plots can also confirm that datasets are properly aligned.

For instance, we can plot *S100A9*, which the Wilcoxon test identified as a marker for *Cluster 1*, and *MS4A1*, a marker for *Cluster 4*:

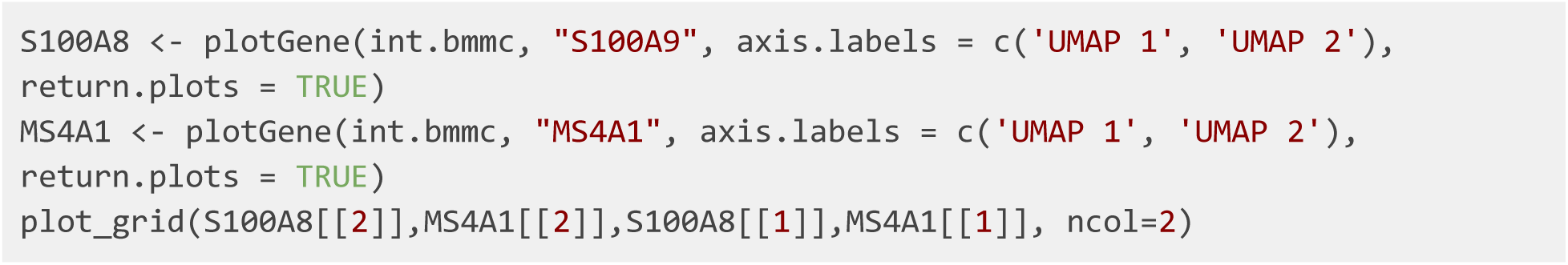

The UMAP plots of expression and chromatin accessibility indicate that *S100A9* and *MS4A1* are indeed specific markers for *Cluster 1* and *Cluster 4*, respectively, with high values in these cell groups and low values elsewhere (**Figure 7**). Furthermore, we can see that the distributions are strikingly similar between the RNA and ATAC datasets, indicating that LIGER has properly integrated the two data types.

**Figure 7:**
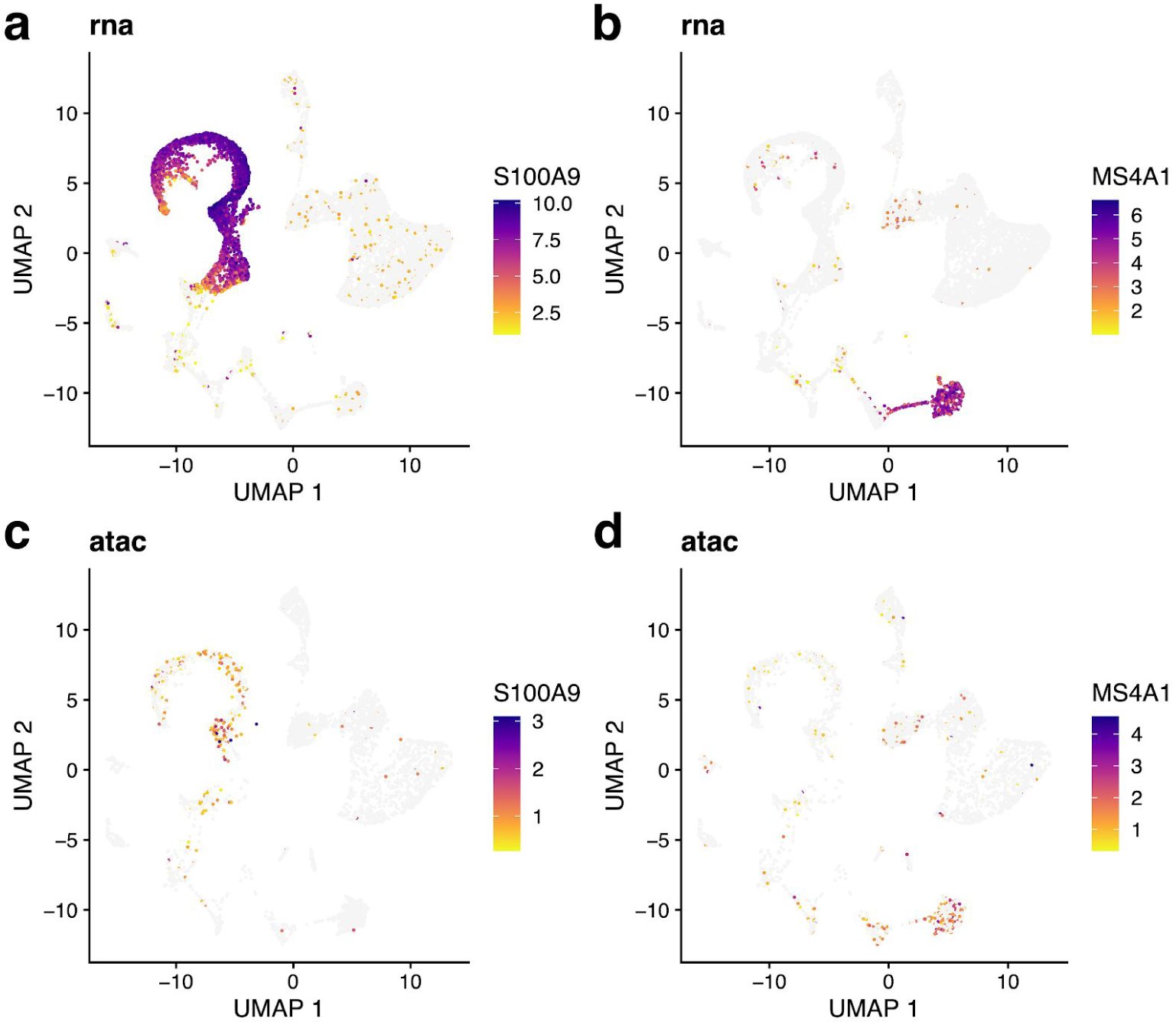
Expression and chromatin accessibility of marker genes selected by LIGER show consistency across modalities. **a,b**, UMAP representations of expression for genes *S100A9* (**a**) and *MS4A1* (**b**). **c,d**, UMAP representations of chromatin accessibility for genes *S100A9* (**c**) and *MS4A1* (**d**), which show highly similar distributions compared to their expression (**a, b**).

**16**. A key advantage of using iNMF instead of other dimensionality reduction approaches such as PCA is that the dimensions are individually interpretable. For example, a single dimension of the space often captures a particular cell type. Furthermore, iNMF identifies both shared and dataset-specific features along each dimension, giving insight into exactly how corresponding cells across datasets are both similar and different. The function *plotGeneLoadings* allows visual exploration of such information. It is recommended to call this function into a PDF file due to the large number of plots produced.

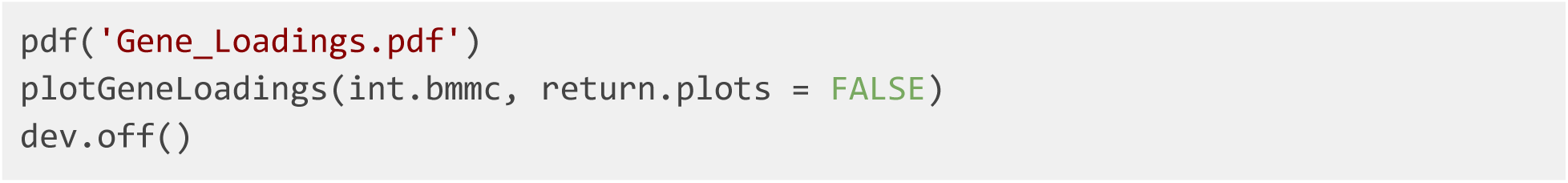

Alternatively, the function can return a list of plots. For example, we can visualize the factor loading of *Factor 7* typing in:

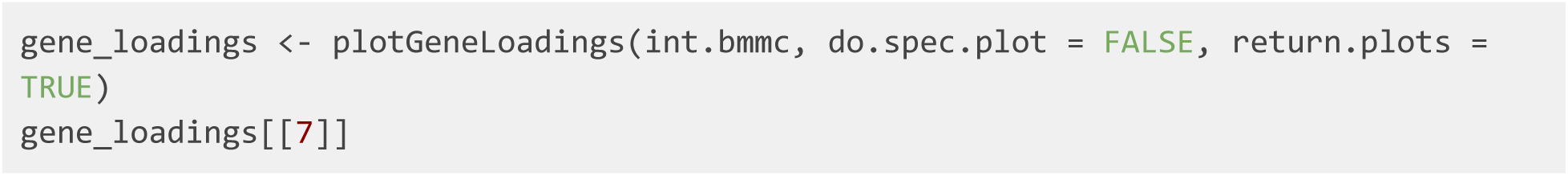

The loading pattern of *Factor 7* shows that *Factor 7* specifically loads on *Cluster 4* (**Figure 8**, top). We also see both the shared markers (including *MS4A1*, which we already inspected above) and dataset-specific genes that characterize this dimension (**Figure 8**, bottom). For example, *CCR6* and *NCF1* are the top dataset-specific genes in the ATAC and RNA datasets, respectively. To inspect these genes, we plotted their expression and accessibility, which confirm that these genes show clear differences (**Figure 9**). *CCR6* shows nearly ubiquitous chromatin accessibility but is expressed only in clusters 2 and 4. The accessibility is highest in these clusters, but the ubiquitous accessibility suggests that the expression of *CCR6* is somewhat decoupled from its accessibility, likely regulated by other factors. Conversely, *NCF1* shows high expression in clusters 1, 3, 4 and 9, despite no clear enrichment in chromatin accessibility within these clusters 4 and 9. This may again indicate decoupling between the expression and chromatin accessibility of *NCF1*. Another possibility is that the difference is due to technical effects--the gene body of *NCF1* is short (∼15KB), and short genes are more difficult to capture in snATAC-seq than in scRNA-seq because there are few sites for the ATAC-seq transposon to insert.

**Figure 8:**
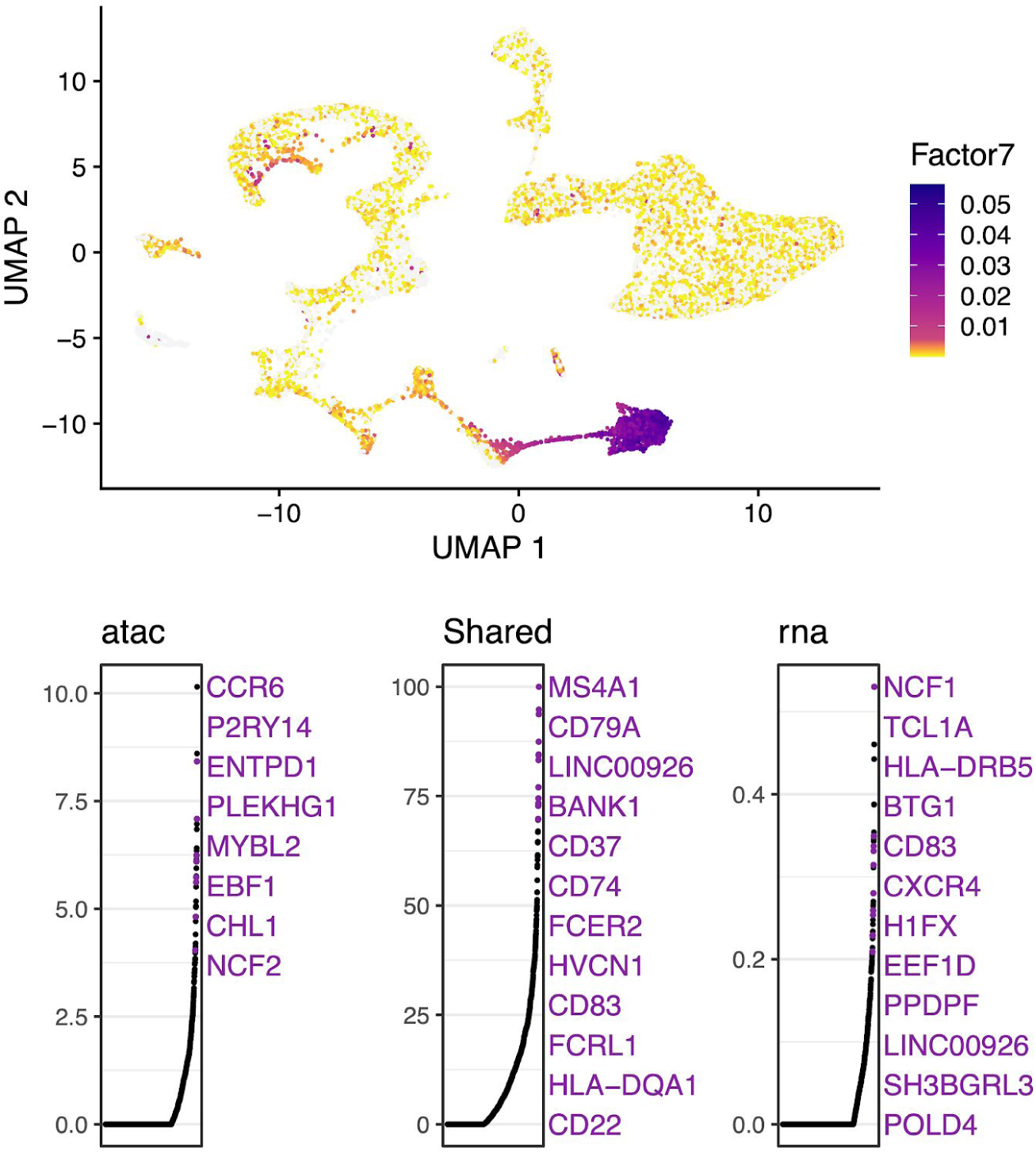
Metagenes and metagene expression levels for BMMC data. UMAP plots showing metagene expression levels (top) and gene loading values (bottom) for Factor 7, which specifically loads on Cluster 4. In gene loading plots, gene names are sorted in decreasing order of magnitude of their factor loading contribution and correspond to colored points in scatterplots. Plots are organized to show the metagene specific to snATAC-seq (left), the shared metagene common to all datasets (middle) and the metagene specific to scRNA-seq profiles (right).

**Figure 9:**
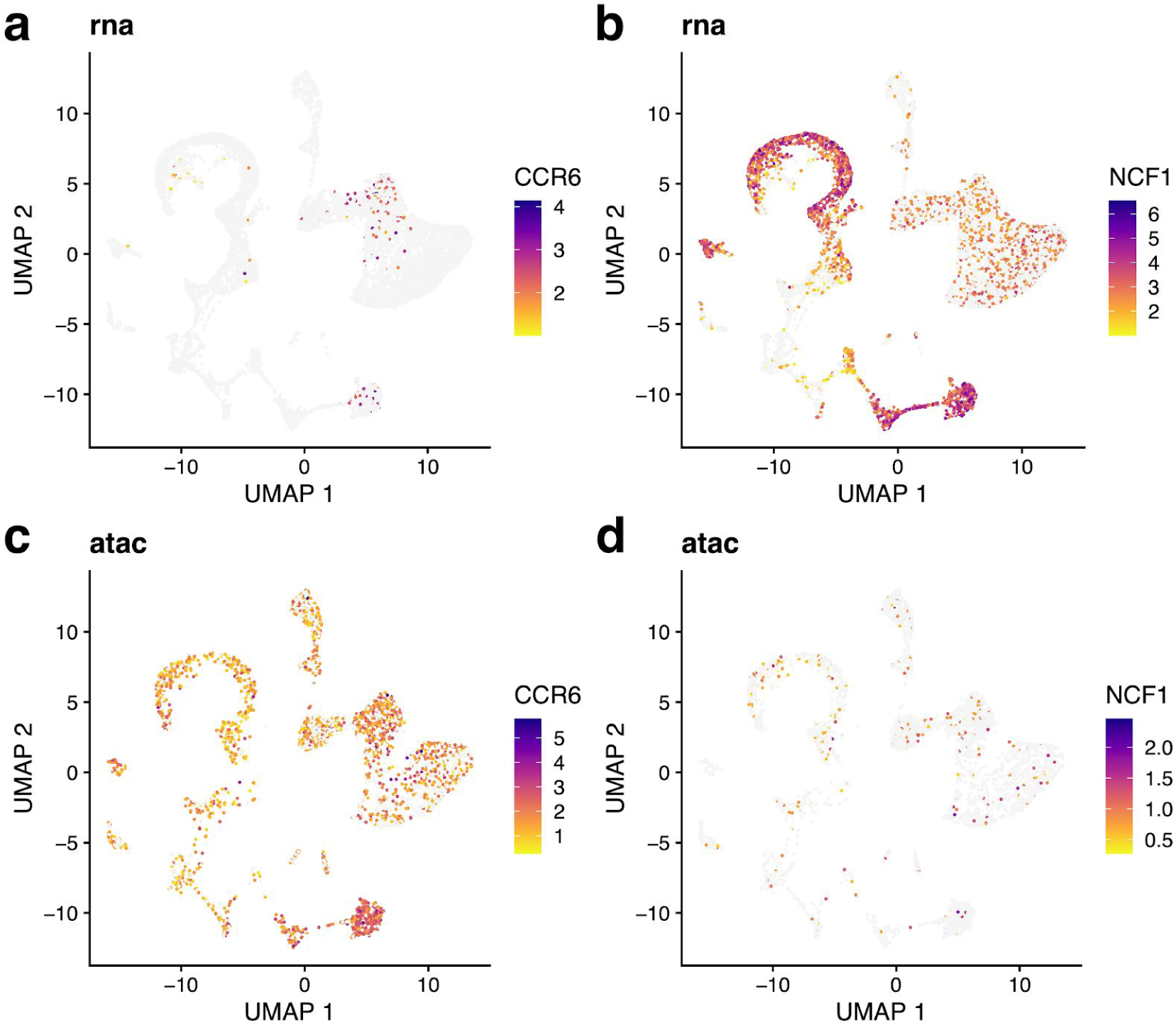
Genes showing expression and accessibility differences. **a,b**, UMAP representation of expression for *CCR6* (**a**) and *NCF1* (**b**). **c,d**, UMAP representation of chromatin accessibility of *CCR6* (**c**) and *NCF1* (**d**), which both show distinct distributions compared to their expressions (**a, b**).

**17**. Single-cell measurements of chromatin accessibility and gene expression provide an unprecedented opportunity to investigate epigenetic regulation of gene expression. Ideally, such investigation would leverage paired ATAC-seq and RNA-seq from the same cells, but such simultaneous measurements are not generally available. However, using LIGER, it is possible to computationally infer “pseudo-multi-omic” profiles by linking scRNA-seq profiles--using the jointly inferred iNMF factors--to the most similar snATAC-seq profiles. After this imputation step, we can perform downstream analyses as if we had true single-cell multi-omic profiles. For example, we can identify putative enhancers by correlating the expression of a gene with the accessibility of neighboring intergenic peaks across the whole set of single cells.

To achieve this, we first need a matrix of accessibility counts within intergenic peaks. The CellRanger pipeline for snATAC-seq outputs such a matrix by default, so we will use this as our starting point. The count matrix, peak genomic coordinates, and source cell barcodes output by CellRanger are stored in a folder named *filtered_peak_matrix* in the output directory. The user can load these and convert them into a peak-level count matrix by typing these commands:

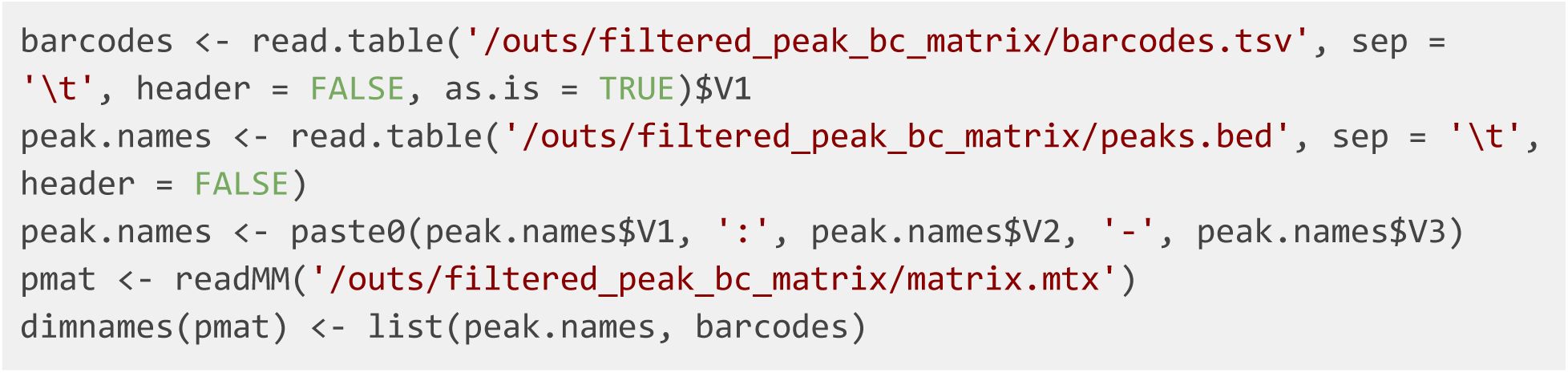

**18**. The peak-level count matrix is usually large, containing hundreds of thousands of peaks. We next filter this set of peaks to identify those showing cell-type-specific accessibility. To do this, we perform the Wilcoxon rank-sum test and pick those peaks which are differentially accessible within a specific cluster. Before running the test, however, we need to: (1) subset the peak-level count matrix to include the same cells as the gene-level counts matrix; (2) replace the original gene-level counts matrix in the LIGER object by peak-level counts matrix; and (3) normalize peak counts to sum to 1 within each cell. We can do this with the following steps:

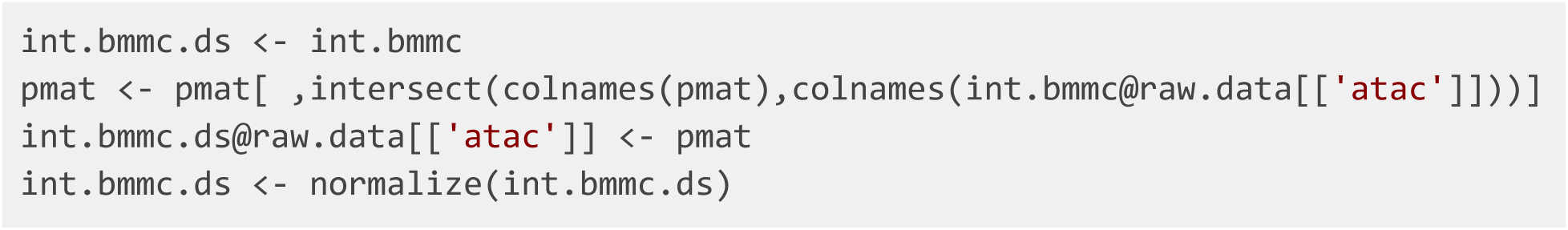

Now we can perform the Wilcoxon test:

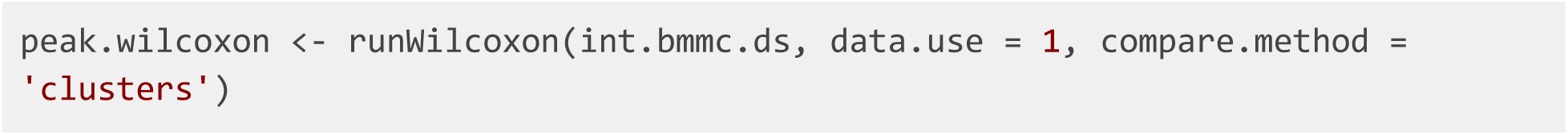

The user can find documentation of important parameters of *runWilcoxon* in the section above (“**Identify Gene Markers of Individual Cell Types**”).

**?TROUBLESHOOTING**

**19**. We can now use the results of the Wilcoxon test to retain only peaks showing differential accessibility across our set of joint clusters. Here we kept peaks with Benjamini-Hochberg adjusted p-value < 0.05 and log fold change > 2.

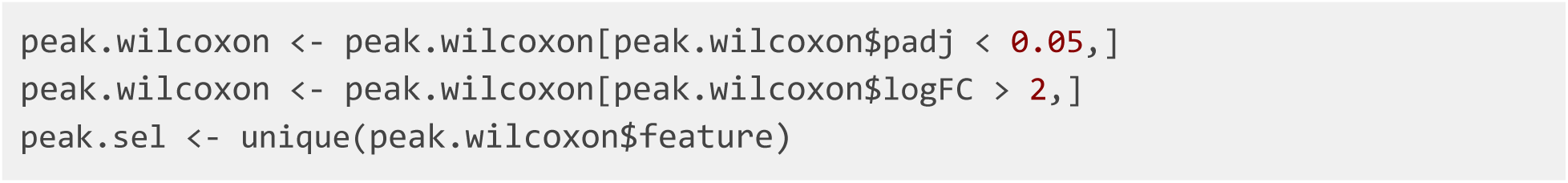

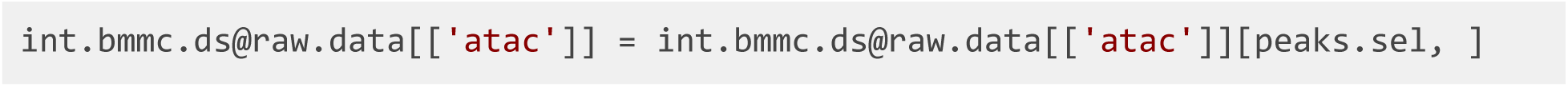

**20**. Using this set of differentially accessible peaks, we now impute a set of “pseudo-multi-omic” profiles by inferring the intergenic peak accessibility for scRNA-seq profiles based on their nearest neighbors in the joint LIGER space. LIGER provides a function named *imputeKNN* that performs this task, yielding a set of profiles containing both gene expression and chromatin accessibility measurements for the same single cells:

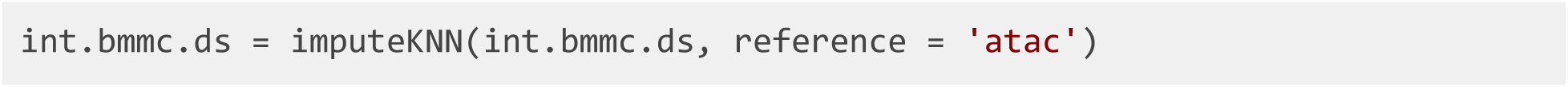

Important parameters of *imputeKNN* are as follows:

- *reference*. Dataset containing values to impute into query dataset(s). For example, setting *reference = ‘atac’* uses the values in dataset *‘atac’* to predict chromatin accessibility values for scRNA-seq profiles.
- *queries*. Dataset to be augmented by imputation. For example, setting *query = ‘rna’* predicts chromatin accessibility values for scRNA-seq profiles.
- *knn_k*. The maximum number of nearest neighbors to use for imputation. The imputation algorithm simply builds a k-nearest neighbor graph using the aligned LIGER latent space, then averages values from the reference dataset across neighboring cells. The default value is 20.
- *weight*. This indicates whether to use KNN distances to weight datasets (TRUE) or to average equally among all neighbors (FALSE). The default is TRUE.
- *norm*. This indicates whether to normalize data after imputation. The default is TRUE.
- *scale*. This indicates whether to scale data after imputation. The default is FALSE.

**21**. Now that we have both the (imputed) peak-level counts matrix and the (observed) gene expression counts matrix for the same cells, we can evaluate the relationships between pairs of genes and peaks, linking genes to putative regulatory elements. We use a simple strategy to identify such gene-peak links: Calculate correlation between gene expression and peak accessibility of all peaks within 500 KB of a gene, then retain all peaks showing statistically significant correlation with the gene. The *linkGenesAndPeaks* function performs this analysis:

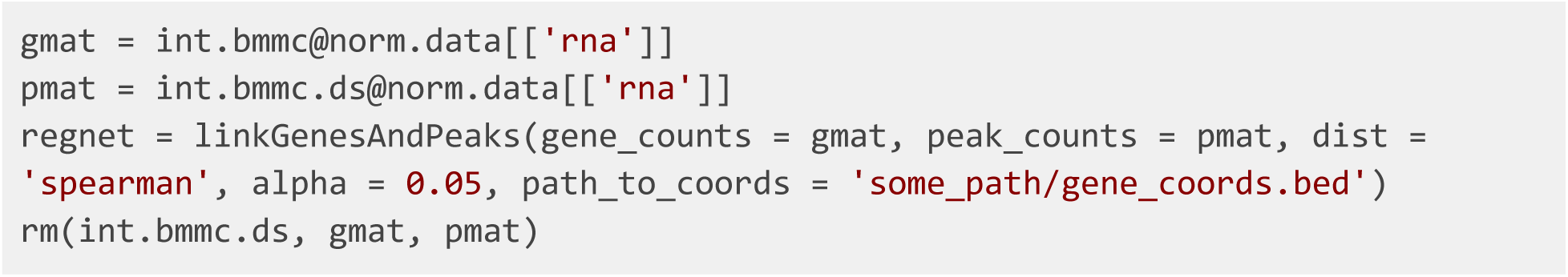

Important parameters of *linkGenesAndPeaks* are as follows:

- *gene_counts*. A gene expression matrix (genes by cells) of normalized counts. This matrix has to share the same column names (cell barcodes) as the matrix passed to *peak_counts*.
- *peak_counts*. A peak-level matrix (peaks by cells) of normalized accessibility values, such as the one resulting from *imputeKNN*. This matrix must share the same column names (cell barcodes) as the matrix passed to *gene_counts*.
- *genes.list*. A list of the genes symbols to be tested. If not specified, this function will use all the gene symbols from the matrix passed to *gmat* by default.
- *dist*. This indicates the type of correlation to calculate -- one of “spearman” (default), “pearson”, or “kendall”.
- *alpha*. Significance threshold for correlation p-value. Peak-gene correlations with p-values below this threshold are considered significant. The default is 0.05.
- *path_to_coords*. The path to the gene coordinates file (in .*bed* format). We recommend passing in the same *bed* file used for making barcodes list in **Step 1**.

**22**. The output of this function is a sparse matrix with peak names as rows and gene symbols as columns, with each element indicating the correlation between peak *i* and gene *j* (or 0 if the gene and peak are not significantly linked). For example, we can subset the results for marker gene *S100A9*, which is a marker gene of *Cluster 1* identified in the previous section, and rank these peaks by their correlation:

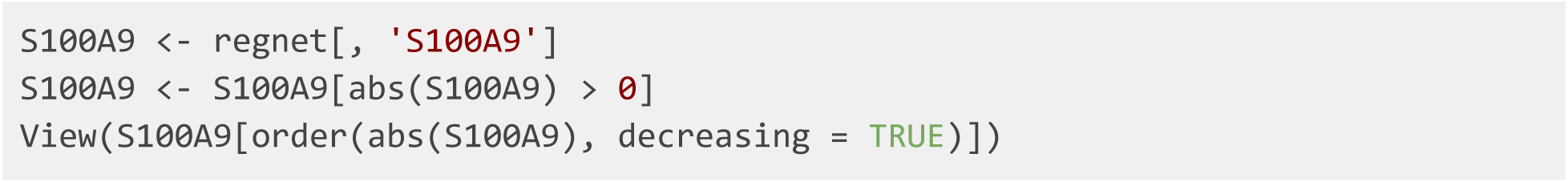

We also provide a function to transform the peaks-genes correlation matrix into an Interact Track for visualizing the calculated linkage between genes and correlated peaks.

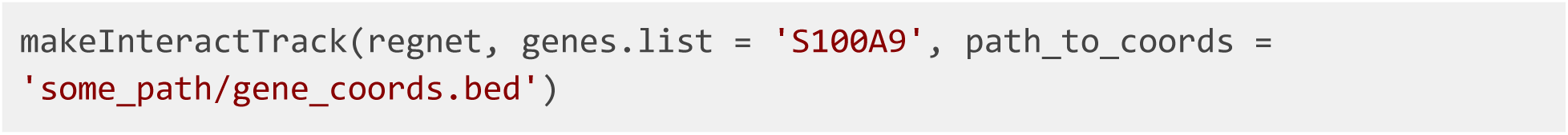

Important parameters of this function are as follows:

- *corr.mat*. A peaks x genes sparse matrix containing inferred gene-peak links (as output by *linkGenesAndPeaks*).
- *genes.list*. A vector of the gene symbols to be included in the output Interact Track file. If not specified, this function will use all the gene symbols in *corr.mat* by default.
- *path_to_coords*. The path to the gene coordinates file (in .*bed* format). We recommend using the same .*bed* file used for making the barcodes list in **Step 1**.
- *output_path*. The path to the directory in which the Interact Track file will be stored. The default is the working directory.

The output of this function will be a UCSC Interact Track file named *‘Interact_Track.bed’* containing linkage information of the specified genes and correlated peaks stored in the given directory. The user can then upload this file as a custom track using this page (https://genome.ucsc.edu/cgi-bin/hgCustom) and display it in the UCSC Genome browser.

For example, the two peaks most correlated to *S100A9* expression are shown in the UCSC genome browser (**Figure 10**). One of the peaks overlaps with the TSS of *S100A8*, a neighboring gene that is co-expressed with *S100A9*, while another peak overlaps with the TSS of *S100A9* itself. The last peak, *chr1:153358896-153359396*, does not overlap with a gene body and shows strong H3K27 acetylation across ENCODE cell lines, indicating that this is likely an intergenic regulatory element.

**Figure 10:**
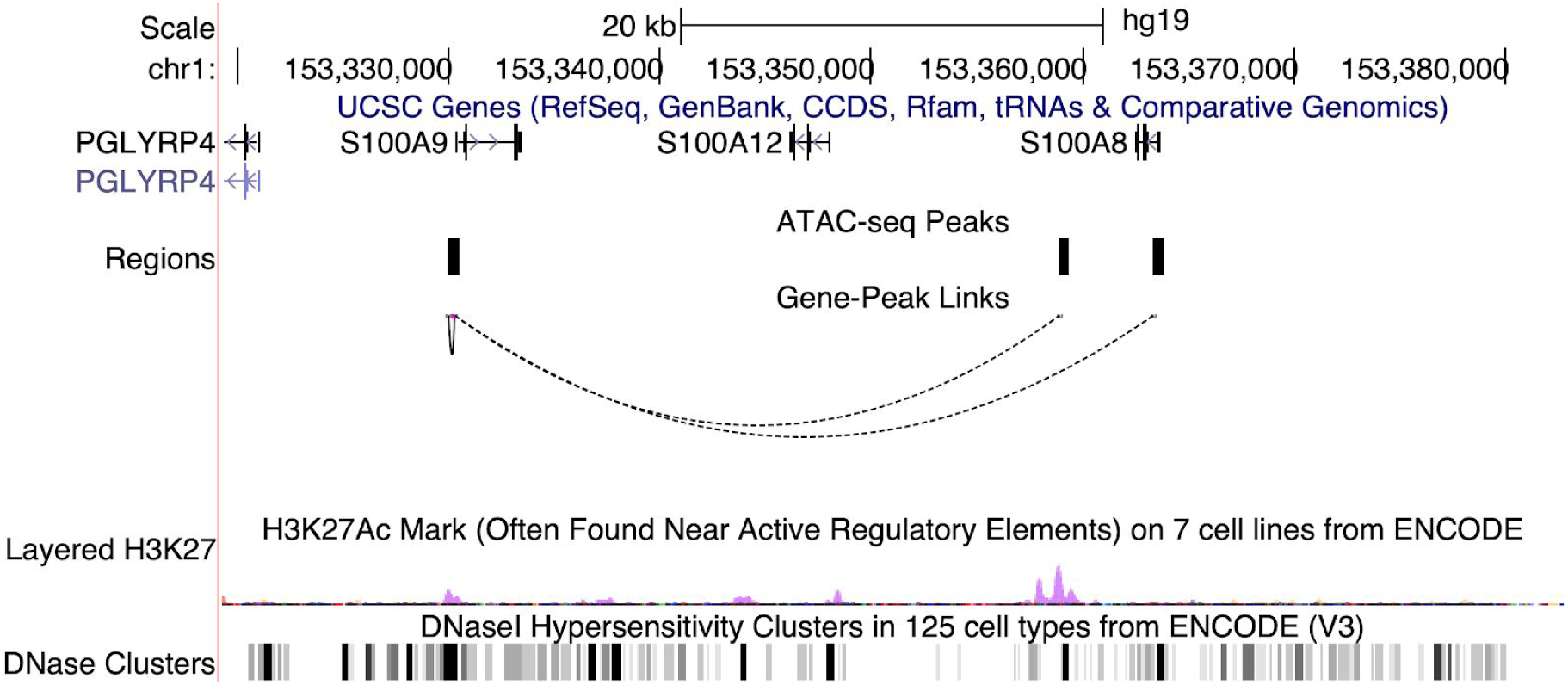
UCSC genome browser view showing the correlations between three candidate chromatin accessible regions and target gene *S100A9*. The locations of three peaks are shown as short black strips within the row “Regions”, and the correlations are illustrated by dotted arcs. H3K27 acetylation and DNasel hypersensitivity across ENCODE cell lines are also shown at the bottom.

To further inspect the correlation between *chr1:153358896-153359396* and *S100A9*, we plotted the accessibility of this peak and the expression of *S100A9* (**Figure 11**). We can see that the two are indeed very correlated and show strong enrichment in clusters 1 and 3. Thus, the intergenic peak likely serves as a cell-type-specific regulator of *S100A9*.

**Figure 11:**
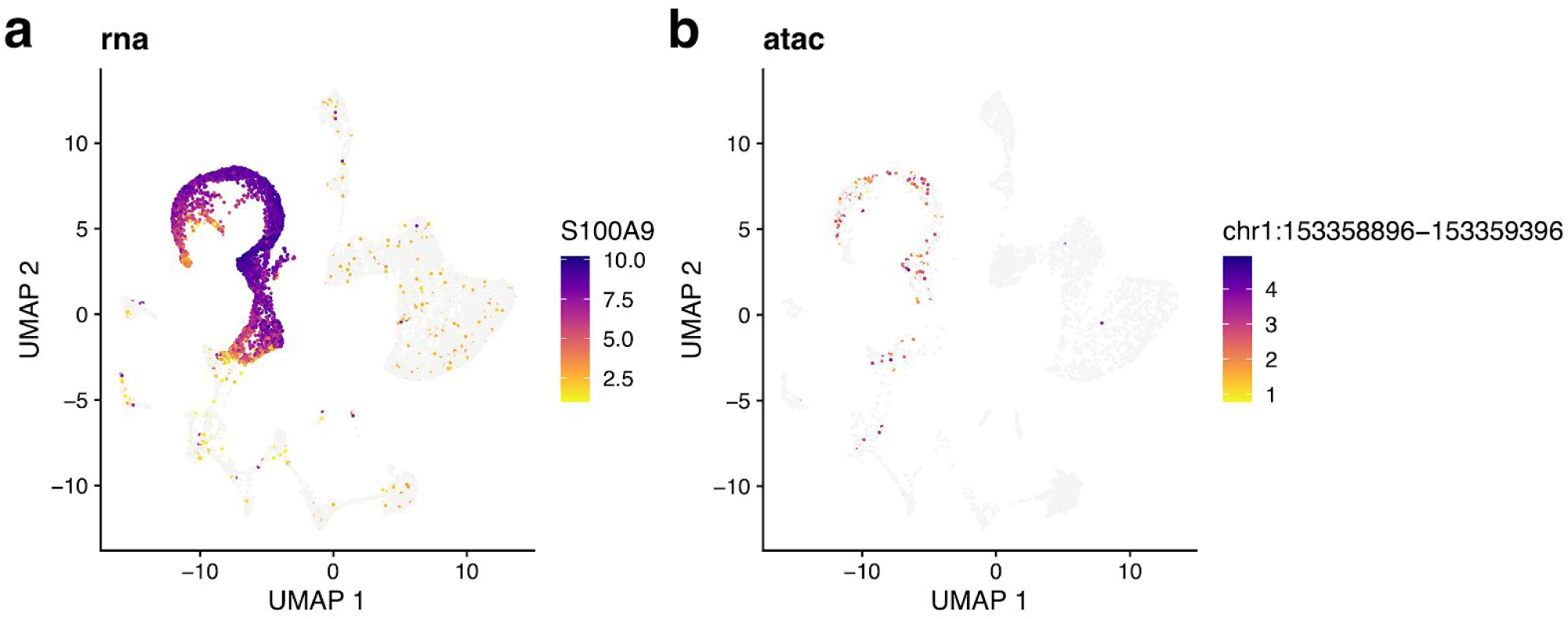
Expression and correlated accessibility for *S100A9* and nearby intergenic peak. **a**, UMAP representation of imputed chromatin accessibility of gene *S100A9*. **b**, UMAP representation of chromatin accessibility for peak *chr1:153358896-153359396.*

### Anticipated Results and Troubleshooting

In this section, we will introduce a new, pre-factorized object to demonstrate several common issues encountered with LIGER and compare possible outputs. This object is composed of two datasets of interneurons and oligodendrocytes from the mouse frontal cortex (Saunders et al. 2018), two distinct cell types that should not align if integrated. We used this dataset in our previous paper as a “negative control” to test whether LIGER spuriously aligns distinct cell types, and we use it here to demonstrate several pitfalls in LIGER analysis.

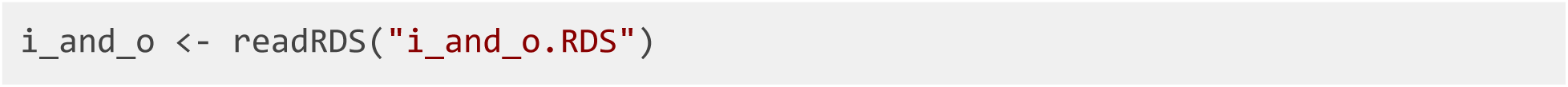

### Selecting hyperparameters

To get the best results from the factorization, we first run a hyperparameter optimization for *k*, the number of factors, and *lambda*, the penalty term associated with dataset specific factors. Although *suggestK* and *suggestLambda* could be used to initially find these values, utilizing *suggestNewK* and *suggestNewLambda* instead, respectively, after running an initial factorization will result in faster output.

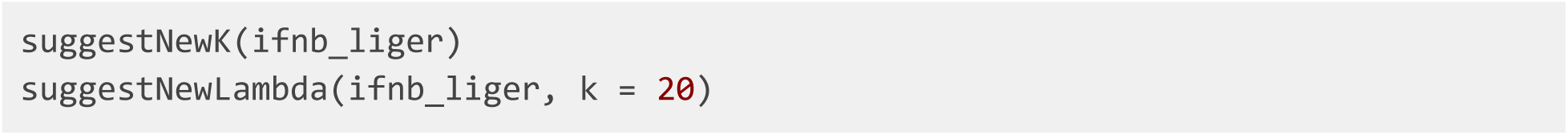

We select the value *k* = 20 at which the increase in median KL divergence becomes negligible, using the plot generated by *suggestNewK* (**Figure 12, b**). The plot generated by *suggestNewLamda* demonstrates that maximum alignment is reached at small values of lambda, so the default value *lambda=5* or less is a reasonable choice for this dataset (**Figure 12, a**). With these parameters, we run *optimizeALS*, LIGER’s implementation of integrative non-negative matrix factorization algorithm, again.

**Figure 12:**
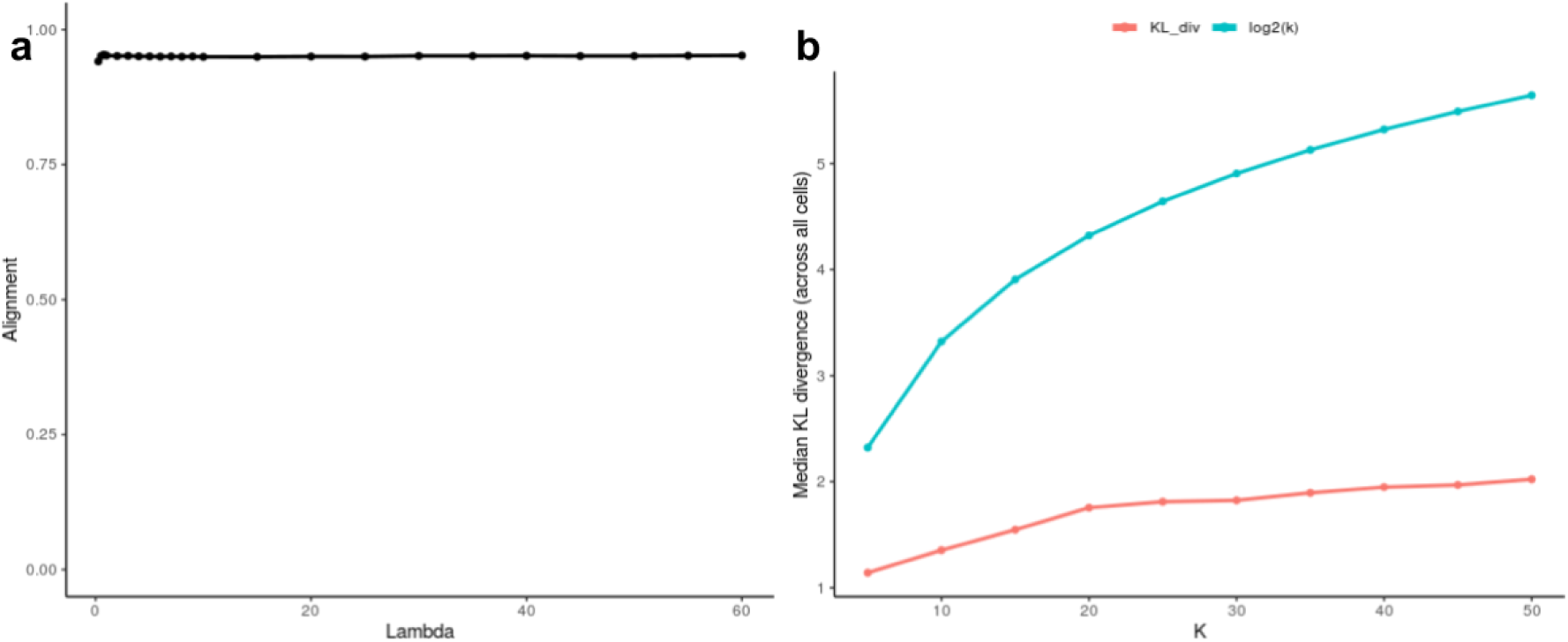
Parameter selection of the number of factors *k* and the tuning parameter *λ*. **a,b**, As increases in *λ* (**a**) and *k* (**b**) results in smaller relative increases in metrics of their effectiveness, the “elbow” of the graph can be interpreted as the optimal hyperparameter value.

We note again that because LIGER is an unsupervised method and no ground truth is available, there is no one correct value of *k*. Thus, we recommend using the above heuristics as a guide rather than a definitive answer. Also, we recommend starting with a value of *k* in the range 20-40 and simply running an initial analysis, rather than trying to determine the perfect *k* before looking at the results.

### Factor Curation

One benefit of iNMF over other dimensionality reduction techniques is the interpretability of the resulting metagenes in terms of biological or technical signals. By studying the gene loadings for each metagene, as represented by the W matrix, we can directly interpret the biological relevance of each factor and exclude nuisance technical or biological signals in downstream analyses. We can also gain insights into the biological interpretation of each factor. *runGSEA* can be used as a tool to analyze factor composition. If no parameters specifying gene sets are given, then the function will use all Reactome gene sets that contain at least one of the genes in the object. This allows a principled means of determining what biological or technical signal each factor represents.

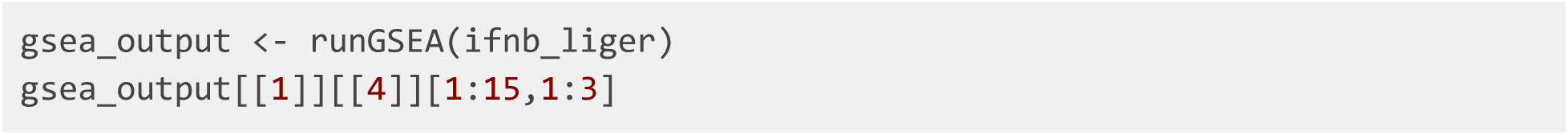

From **Table 3**, we find overrepresentation of gene sets related to the interaction between the innate and adaptive immune systems in Factor 4. The ER-Phagosome pathway is responsible for the release of cytokines as a part of the process of cross presentation. This supports the inclusion of the gene sets for cytokine signalling and interleukins, a type of cytokine. We also see that GPCR ligand binding is significant, possibly meaning that the immune response is a result of non-immune signalling. From this information, we can hypothesize that cells that highly load factor 4 may be responsible for initiating an immune response, specifically that of a T lymphocyte due to the significance of interleukin signalling and cross presentation. Because we know factor 4 loads primarily on cluster 5, the cluster may represent T lymphocytes.

**Table 3:** Gene sets enriched in factor 16 of IFNB LIGER result.

|  | pathway | pval | padj |
| --- | --- | --- | --- |
| 1 | Immune System | 0.000100 | 0.007639 |
| 2 | Innate Immune System | 0.000100 | 0.007639 |
| 3 | Immunoregulatory interactions between a Lymphoid and a non-Lymphoid cell | 0.000100 | 0.007639 |
| 4 | Antigen processing-Cross presentation | 0.000100 | 0.007639 |
| 5 | Adaptive Immune System | 0.000200 | 0.007639 |
| 6 | Cytokine Signaling in Immune system | 0.000200 | 0.007639 |
| 7 | Signaling by Interleukins | 0.000200 | 0.007639 |
| 8 | GPCR ligand binding | 0.000200 | 0.007639 |
| 9 | ER-Phagosome pathway | 0.000201 | 0.007639 |
| 10 | TNFR2 non-canonical NF-kB pathway | 0.000402 | 0.013750 |

**Table 4:**
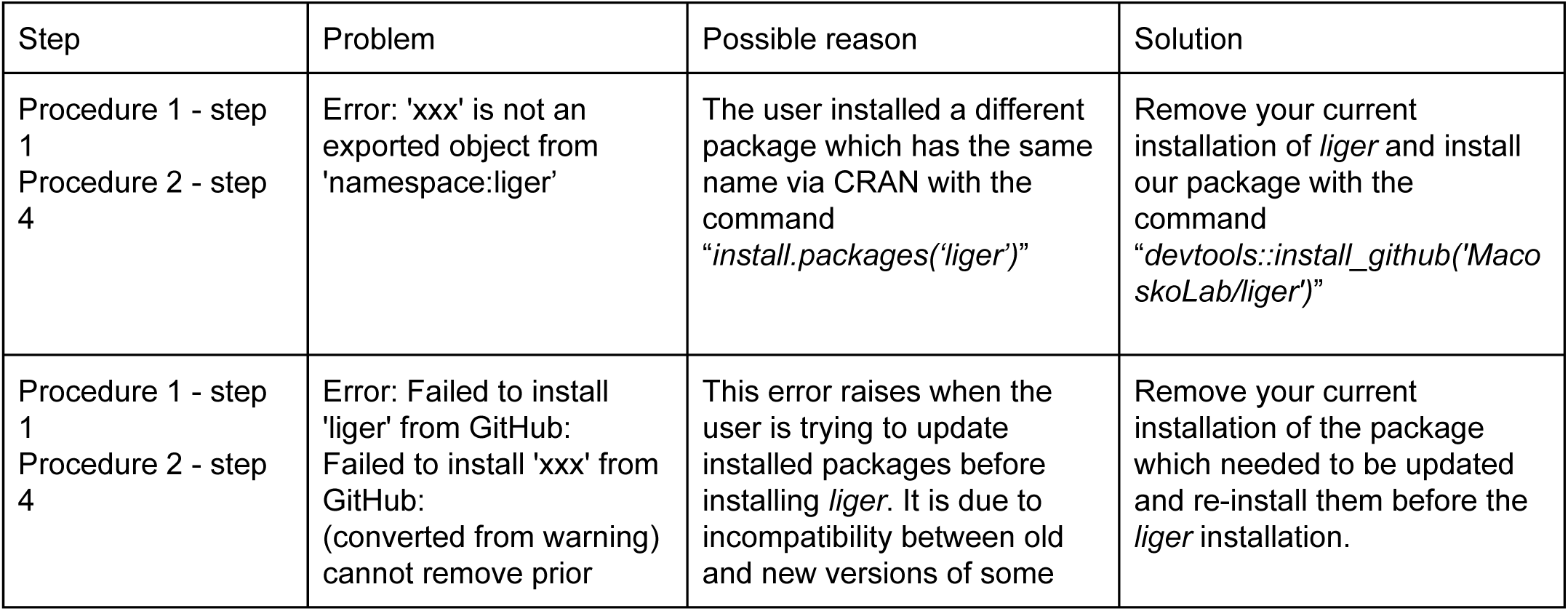
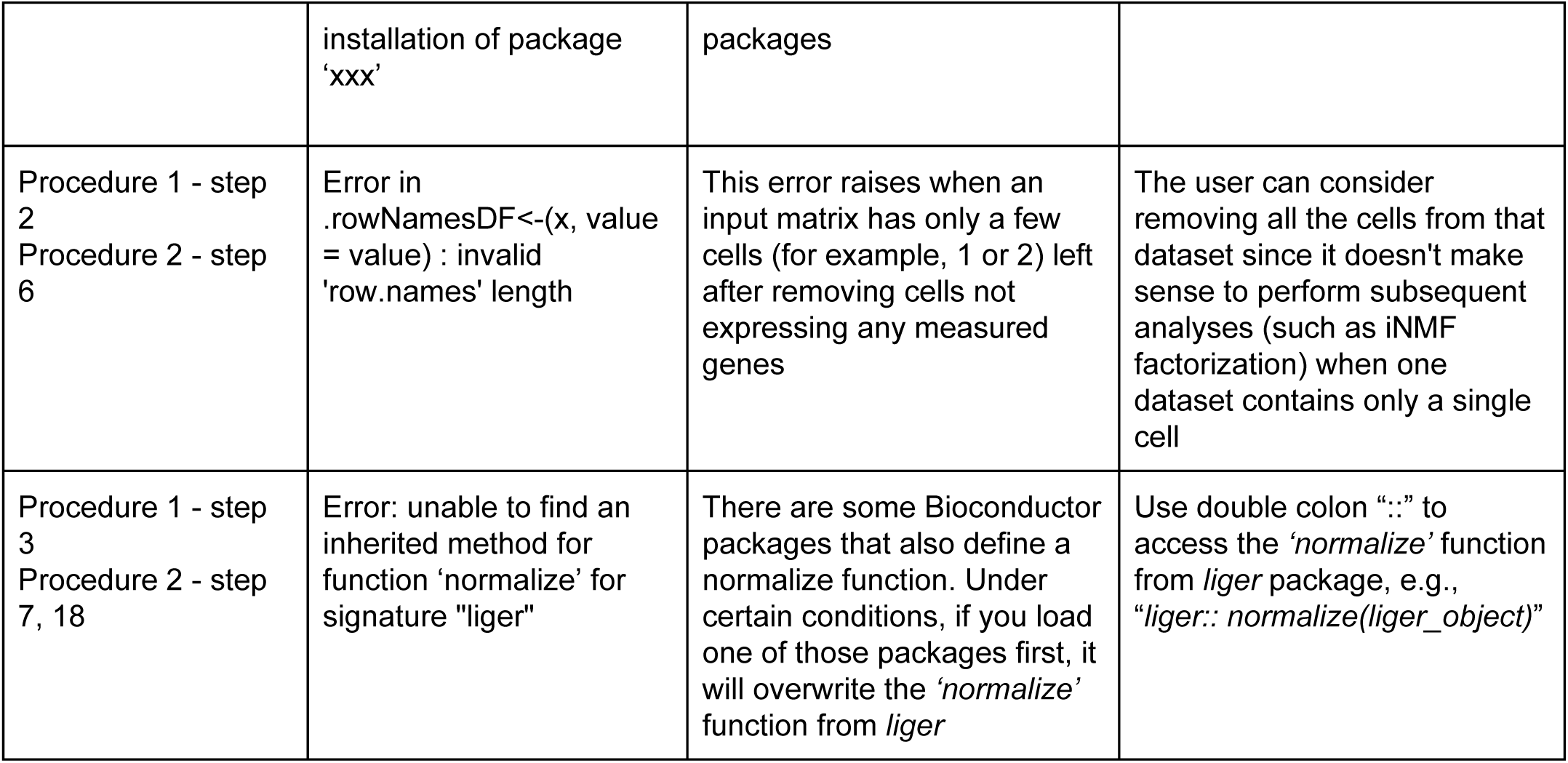
Troubleshooting table.

If we have a custom list of gene sets to study, we can pass those to *runGSEA* in the form of a named list of Entrez IDs. This input can easily be generated with the help of the *msigdbr* package. In the example below, a set of mitochondrial gene sets from the Gene Ontology cellular components subcategory is used for GSEA.

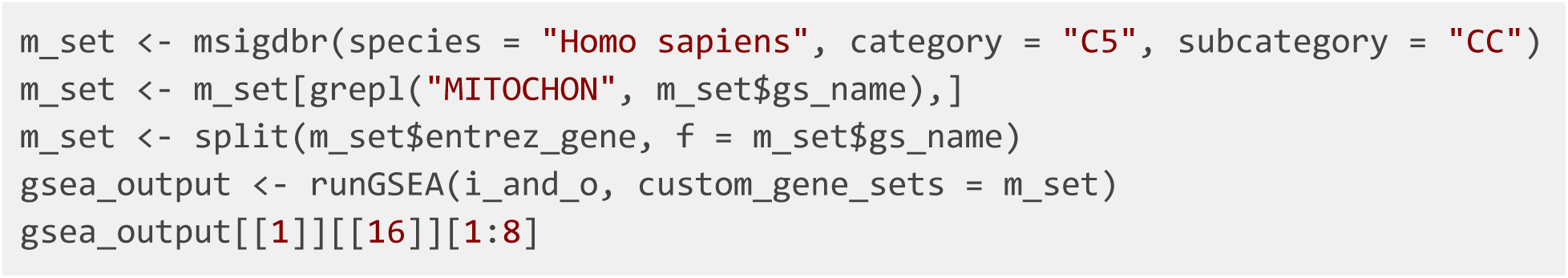

After running GSEA on *i_and_o*’s factors, we find that factors 15 and 16 significantly overrepresent several gene sets associated with mitochondrial function.

We can also use *plotFactors* after running *quantile_norm* to directly compare raw and normalized gene loadings across the datasets. Because of the large number of charts it generates, it should be called into a PDF.

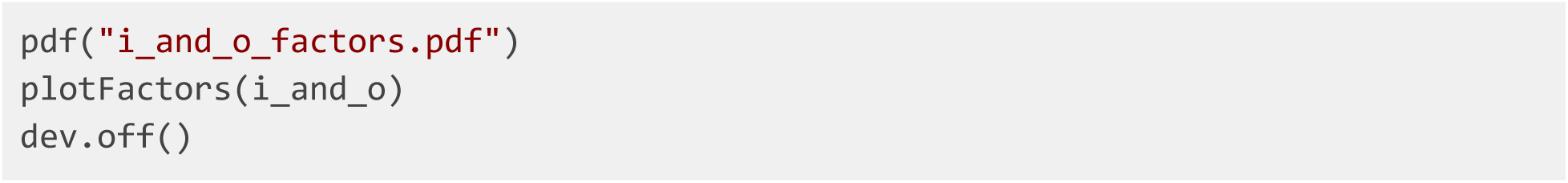

If we look at factor 15, which overexpresses mitochondrial gene sets, we can see that a majority of cells in both datasets have non-zero cell loading values on this factor. This is a common pattern in metagenes representing technical artifacts; biological signals often have much more specific and sparse loadings. Thus, it is likely that this factor captures a technical artifact related to mitochondrial genes and should be removed from the analysis.

To remove these factors from further analysis, we again run *quantile_norm*, with the *dims.use* parameter equal to the set difference of the list of all factors and technical artifacts.

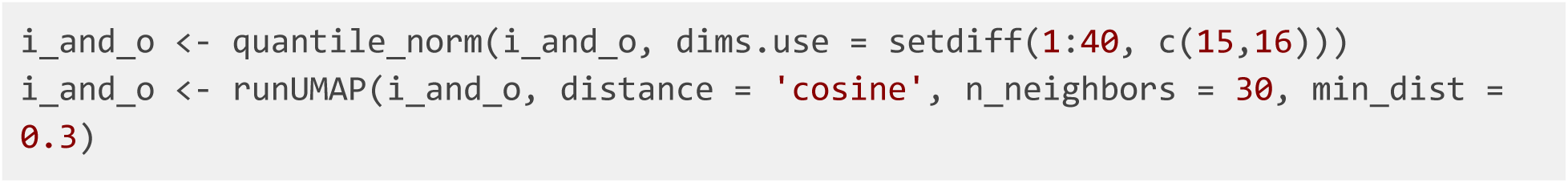

If we compare the final integration with and without the mitochondrial artifact factors, we find that the alignment of the datasets decreases slightly after we remove the factors (**Figure 14**). This is because, without the artifacts, there is less overlap in expression between the datasets for LIGER to use in integration.

**Figure 13:**
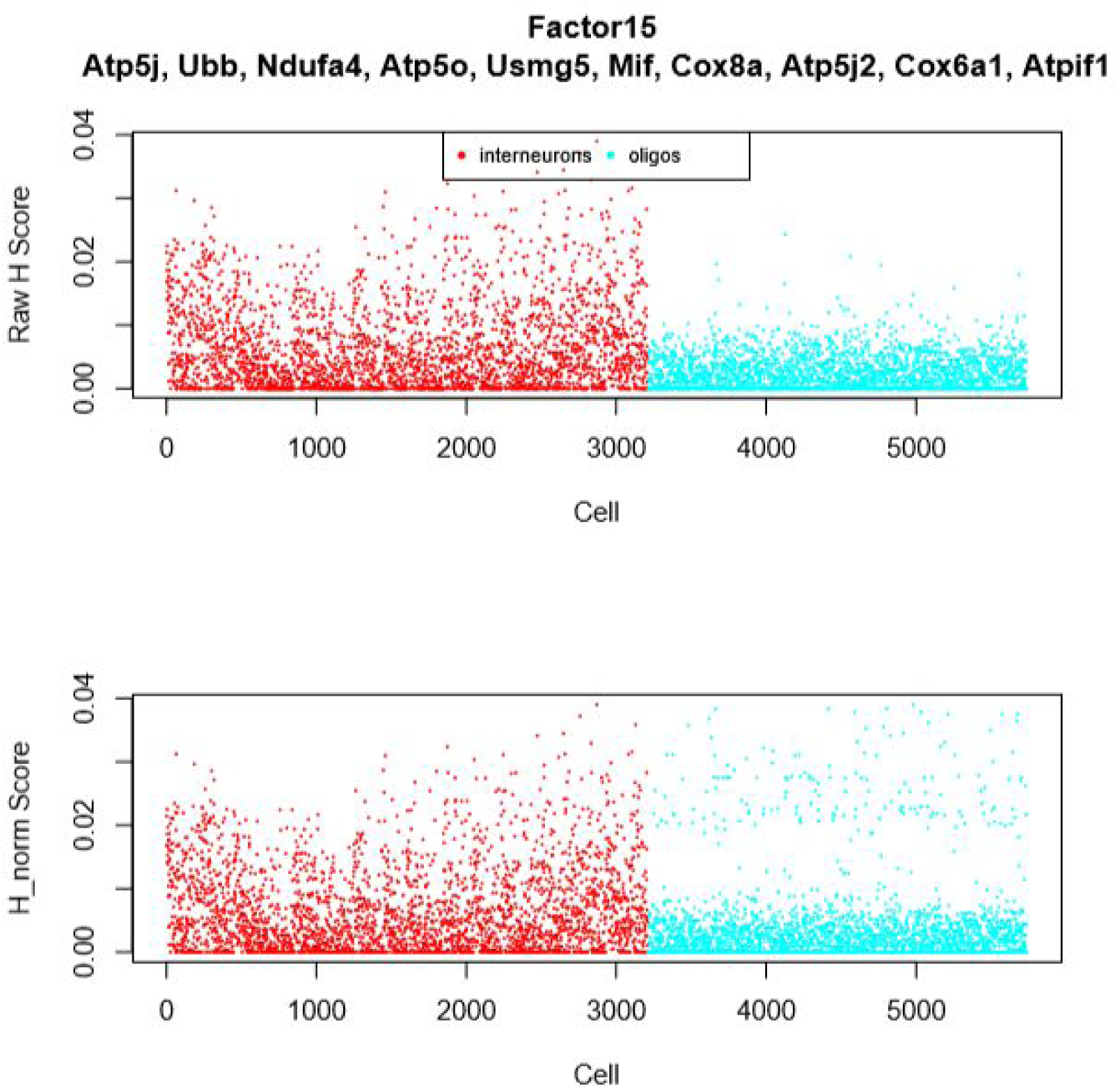
Plots of raw and normalized loading of Factor 15. Scatter plots, with factor loadings values as y axis and cells as x axis, for both unaligned (raw) and aligned (normalized) factor loadings of *Factor 15*.

**Figure 14:**
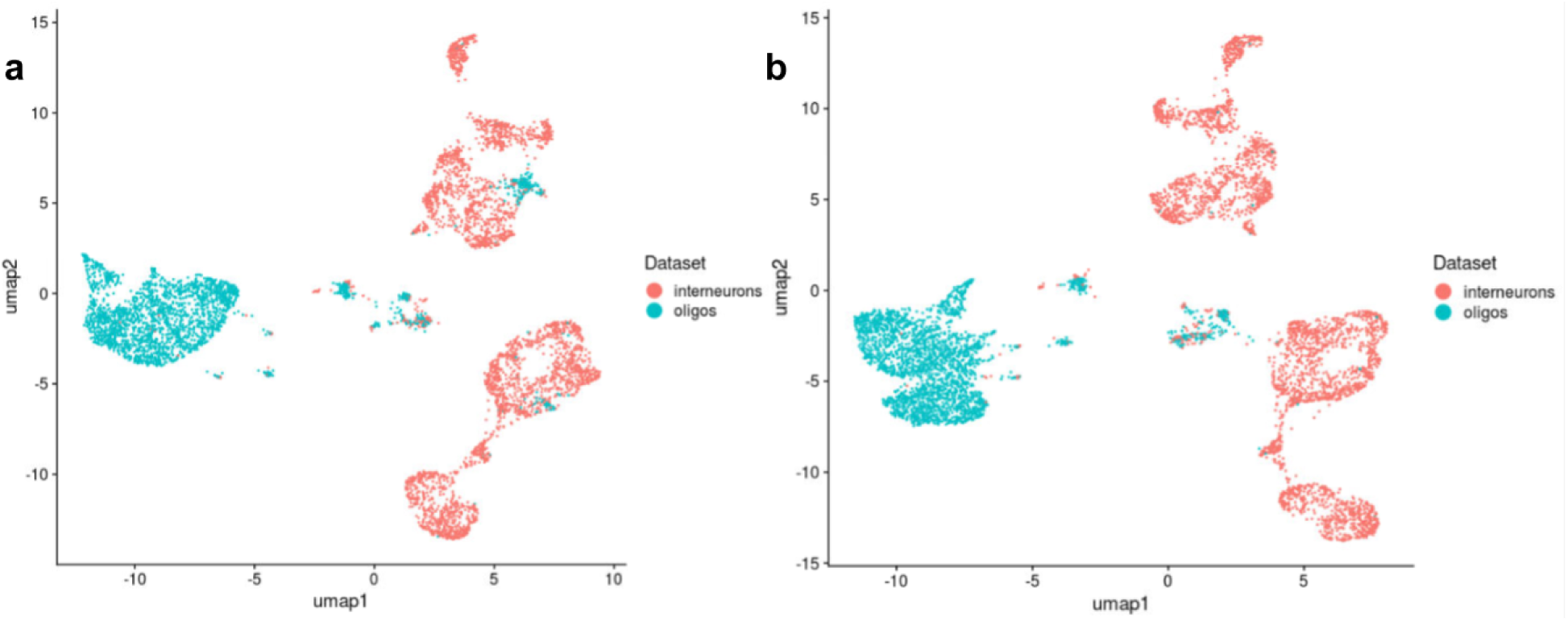
The alignment between datasets decreases after removing the mitochondrial artifact factors. **a,b**, UMAP plots of a LIGER analysis of 3212 interneurons and 2524 oligodendrocytes, with (**a**) and without (**b**) factors 15 and 16, colored by datasets.

### Metrics for Confirming Results

*calcARI*, based on the Adjusted Rand Index, and *calcPurity* can be used to compare the clustering generated by LIGER with some other clustering, such as known cell types or clustering as determined by another method. Both return a value between 0 and 1, with 0 representing total disagreement and 1 representing identical clusterings.

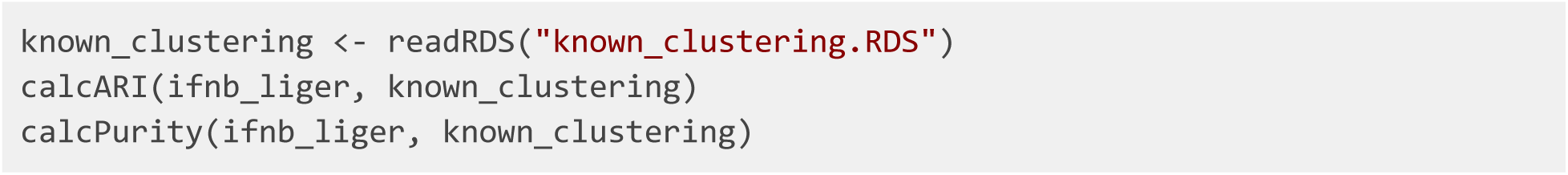

*calcAgreement* returns a metric of the distortion of the geometry of the datasets after factorization and quantile alignment. Although it can theoretically approach a maximum of 1, representing complete preservation of geometry, it generally reaches 0.2 to 0.3.

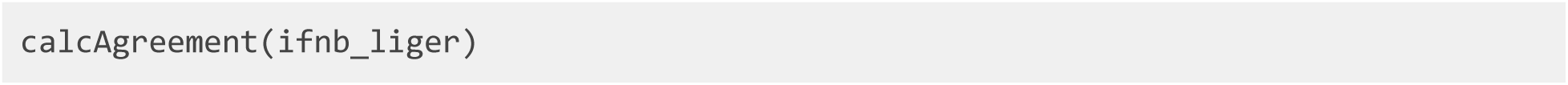

*calcAlignment* returns a metric of how uniformly mixed multiple datasets are, with a maximum of 1 representing perfect integration.

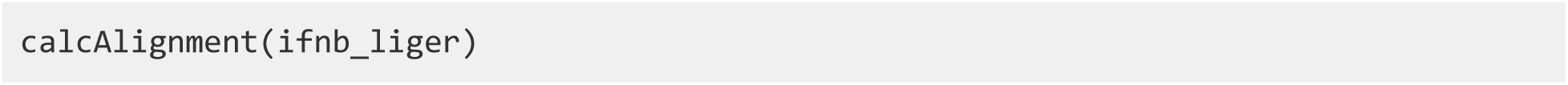

Here we show side-by-side UMAP representations of the oligos/interneurons dataset and the PBMC dataset (**Figure 15**). This plot indicates the poor alignment between the oligos and interneurons visually, in addition to the metrics calculated above.

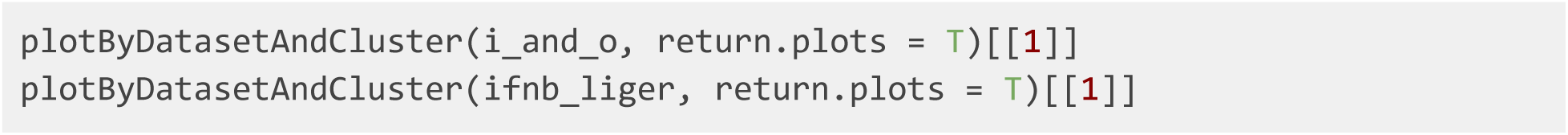

**Figure 15:**
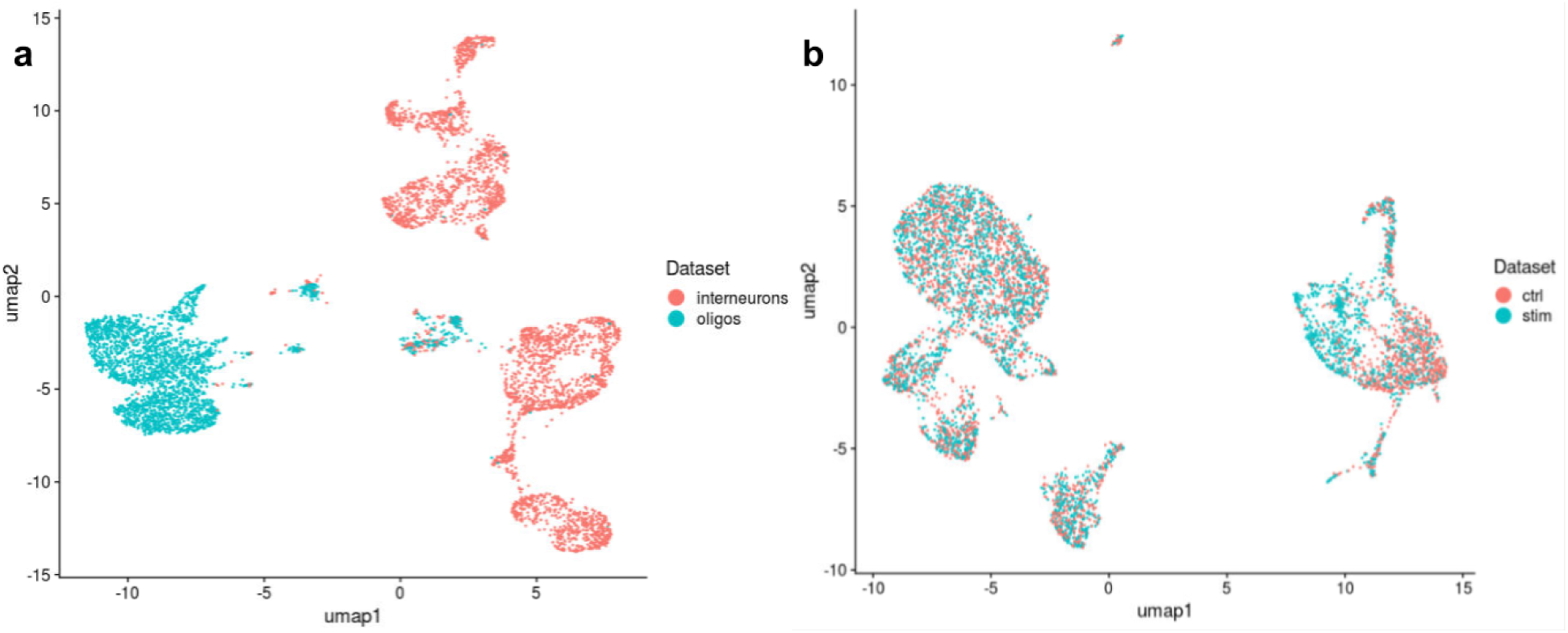
Distinct cell types show poor alignment compared to normal PBMC datasets. **a**, UMAP plot of a LIGER analysis of two distinct cell types, interneurons (3212 cells) and oligos (2524 cells) showing poor alignment. **b**, UMAP plot of a LIGER analysis of 3000 control and 3000 interferon-stimulated PBMCs showing complete and well-mixed alignments.

In this example, we see very limited overlap between the *interneuron* and *oligos* datasets in *i_and_o*, whereas the control and stimulated datasets in *ifnb_liger* are almost perfectly aligned. If you see a plot like the one on the left, it is likely that you are trying to integrate datasets that have no common biology. In addition to visually inspecting the t-SNE or UMAP plot, you can use the function *calcAlignment* to calculate a metric quantifying the degree of alignment.

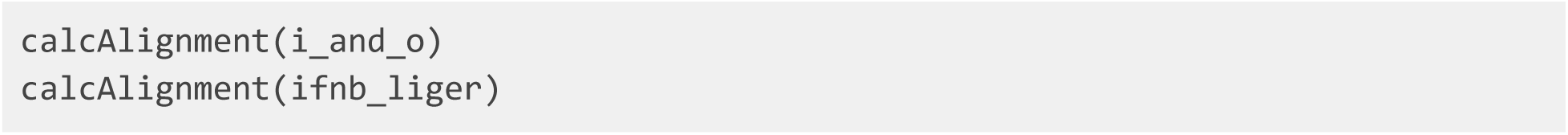

Returning to our example above, the oligodendrocytes and interneurons dataset gives an alignment score of 0.161, whereas *ifnb_liger* has a near-perfect alignment score of 0.947.

## Acknowledgments

This work was supported by NIH grants R01 AI149669-01 and R01 HG010883-01 (JDW) and U19 1U19MH114821 (EZM).

## Author Contributions

CG, JL, JS, and JDW performed the data analysis. CG, JL, JS, and JDW wrote the paper, with input from EZM and VK. All authors read and approved the final manuscript.

